# The Structures of SctK and SctD from *Pseudomonas aeruginosa* Reveal the Interface of the Type III Secretion System Basal Body and Sorting Platform

**DOI:** 10.1101/2020.09.22.308627

**Authors:** Meenakumari Muthuramalingam, Sean K. Whittier, Scott Lovell, Kevin P. Battaile, Shoichi Tachiyama, David K. Johnson, Wendy L. Picking, William D. Picking

## Abstract

Many Gram-negative bacterial pathogens use type III secretion systems (T3SS) to inject effector proteins into eukaryotic cells to subvert normal cellular functions. The T3SS apparatus (injectisome) shares a common overall architecture in all systems studied thus far, comprising three major components – the cytoplasmic sorting platform, the envelope-spanning basal body and the external needle with controlling tip complex. The sorting platform consists of an ATPase (SctN) connected to “pods” (SctQ) having six-fold symmetry via radial spokes (SctL). These pods interface with the 24-fold symmetric SctD inner membrane ring (IR) via an adaptor protein (SctK). Here we report the first high-resolution structure of a SctK protein family member, PscK from *Pseudomonas aeruginosa*, as well as the structure of its interacting partner, the cytoplasmic domain of PscD (SctD). The cytoplasmic domain of PscD forms a forkhead-associated (FHA) fold, like that of its homologues from other T3SS. PscK, on the other hand, forms a helix-rich structure that does not resemble any known protein fold. Based on these structural findings, we present the first model for an interaction between proteins from the sorting platform and the IR. We also test the importance of the PscD residues predicted to mediate this electrostatic interaction using a two-hybrid analysis. The functional need for Arg96 *in vivo* was then confirmed by monitoring secretion of the effector ExoU. These structures will contribute to the development of atomic-resolution models of the entire sorting platform and to our understanding of the mechanistic interface between the sorting platform and the basal body of the injectisome.

**Highlights:**

- The structures of *Pseudomonas aeruginosa* PscD (SctD) and PscK (SctK) were solved
- The interface between the T3SS basal body and sorting platform was modeled
- The crystal structure of PscK is the first for any SctK family member
- PscK represents a novel protein fold
- Site-directed mutagenesis supports a computational model of the PscD-PscK interface

Graphical abstract.
The first reported structure for a T3SS SctK protein family member was solved for PscK from *Pseudomonas aeruginosa* and this allowed for modeling of the interface between this sorting platform protein and the cytoplasmic domain of PscD (a SctD protein family member). This allowed for identification of amino acid residues that may play a role in the interaction between these proteins. The interface appears to be dominated by electrostatic interactions and mutagenesis confirmed the importance of key residues in driving their interaction based on two-hybrid analysis.

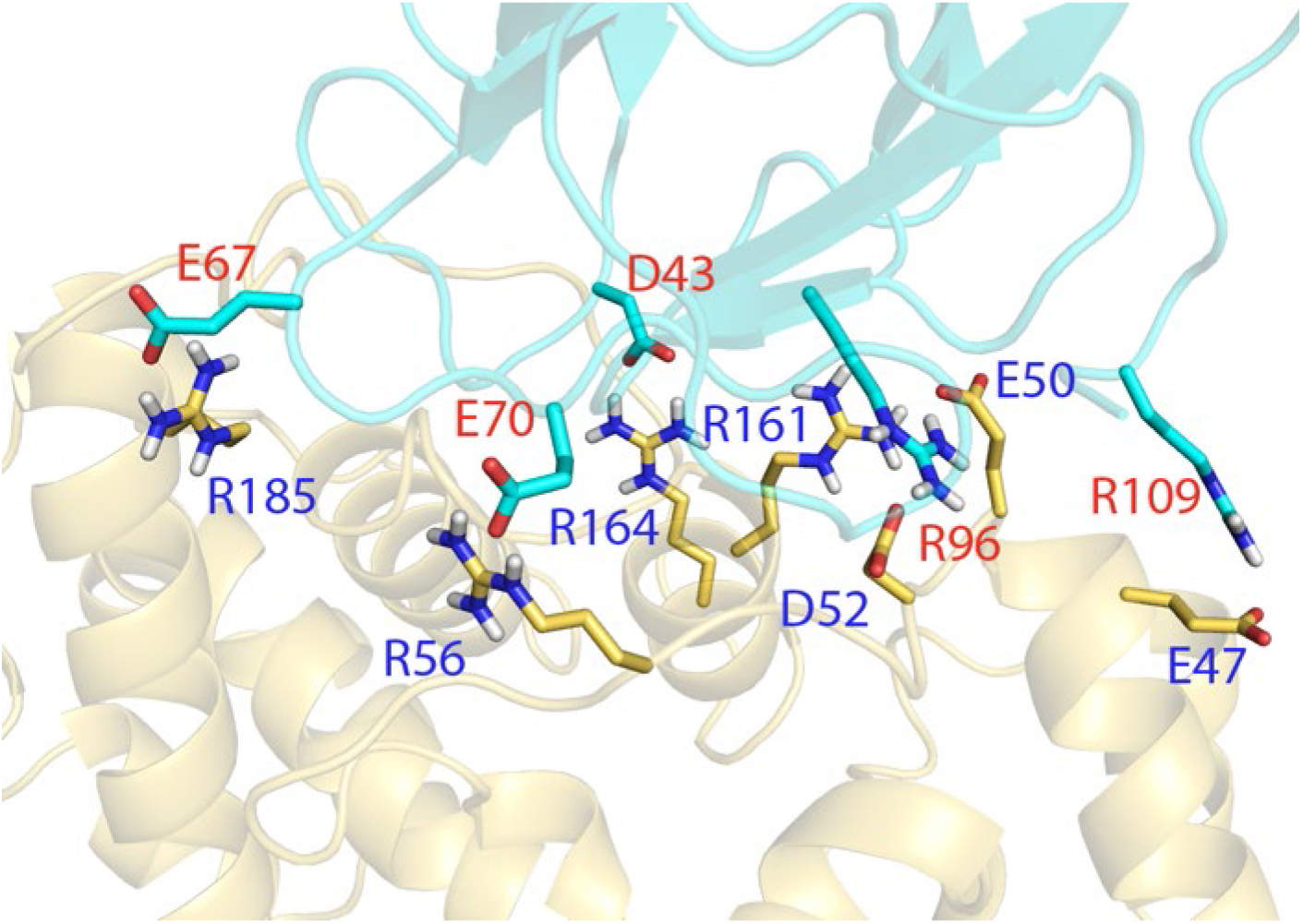

## Introduction

The type III secretion system (T3SS) is used by many Gram-negative pathogens to inject effector proteins from the bacterial cytoplasm into the host cell to promote pathogenesis [1, 2]. T3SS-containing pathogens cause a diverse spectrum of clinical diseases including gastroenteritis and enteric fever (*Salmonella*), dysentery (*Shigella*), plague (*Yersinia*) and whooping cough (*Bordetella*) [3]. While these systems are diverse with regard to their host altering effector proteins, the T3SS apparatus (injectisome) is relatively well conserved with respect to overall structure and architecture. The injectisome is comprised of: 1) a dynamic cytoplasmic sorting platform that is needed for recognizing secretion substrates and controlling/powering the secretion process; 2) an envelope-spanning basal body that is anchored rigidly within the inner membrane, cell wall and outer membrane; and 3) an exposed needle filament with tip complex that extends from the bacterial surface (**Suppl. Fig. S1**). Here we will use the Sct (secretion and cellular translocation) nomenclature for injectisome components except when referring to proteins from specific pathogens (see **Table 1**) [4].

**Table 1.** Major components of the outer membrane ring, inner membrane ring and sorting platform for *Shigella, Salmonella, Yersinia and Pseudomonas aeruginosa*.

| Organism | Sct name | *Shigella* | *Salmonella* | *Yersinia* | *Pseudomonas* |
| --- | --- | --- | --- | --- | --- |
| <u>Basal Body</u> |  |  |  |  |  |
| OM Ring | SctC | MxiD | InvG | YscC | PscC |
| IM Ring | SctD/SctJ | MxiG/MxiJ | PrgH/PrgK | YscD/YscJ | PscD/PscJ |
| Export Gate | SctV | MxiA | InvA | YscV | PscV |
| <u>Sorting Platform</u> |  |  |  |  |  |
| Adaptor | SctK | MxiK | OrgA | YscK | PscK |
| Pods | SctQ | Spa33 | SpaO | YscQ | PscQ |
| Radial spokes | SctL | MxiN | OrgB | YscL | PscL |
| ATPase | SctN | Spa47 | InvC | YscN | PscN |
| Stalk | SctO | Spa13 | InvI | YscO | PscO |

A standard nomenclature has been proposed with the Sct (secretion and cellular translocation) designation used with the letter designation based on the *Yersinia* T3SS gene names.

The injectisome basal body consists of three main structural proteins distributed across the two bacterial membranes. SctD and SctJ make up the inner membrane ring (IR), which has a 24-fold symmetry. Meanwhile, SctC forms the outer membrane ring and has a 15-fold symmetry [5]. The SctD protein family has periplasmic and cytoplasmic domains connected by a single pass transmembrane helix. Beneath the basal body lies the sorting platform, which is formed by a central ATPase (SctN) connected to six radially symmetric SctQ ‘pods’ via SctL proteins. The cytosolic domain of SctD and the SctQ pods of the sorting platform are, in turn, linked through mutual interactions with the adaptor protein SctK [6, 7].

The interaction between the IR and the sorting platform is crucial for active type III secretion. Our understanding of that interaction is limited, in part, because there is no high-resolution structure for any SctK adaptor protein. Here we present the first atomic-resolution structure of the SctK family protein PscK from *Pseudomonas aeruginosa*, together with the structure of its SctD interacting partner in the inner membrane, the cytoplasmic domain of PscD or PscD^C^. We also present a computational model for the PscK-PscD^C^ interface that may be stabilized primarily by electrostatic interactions. We then provide initial experimental support for the importance of at least some electrostatic interactions at the PscD-PscK interface. As key components of a functional injectisome, the information revealed here is expected to help better understand the overall structure and function of the T3SS.

## Results

### The structure of PscK

The T3SS injectisome inner membrane ring consists of the proteins SctD and SctJ having a packed 24-fold symmetry. SctD is a transmembrane protein and its cytoplasmic domain, SctD^C^, forms an interface with the sorting platform protein SctK, which has an evenly spaced six-fold symmetry when bound to the so-called ‘pods’ made up of SctQ. This symmetry change in *Salmonella* is suggested to be accommodated by a reordering of SctD^C^ (PrgH^C^ in *Salmonella*), which is enabled by a linker between it and its transmembrane helix [6, 8](REFS). This does not appear to be the case in *Shigella* where SctD^C^ (MxiG^C^) remains in an evenly spaced 24-fold symmetry configuration within the wild-type injectisome [9]. Thus, there is still much to learn about the architectural differences between even closely related T3SS.

Unlike SctD, for which examples of high resolution structures are available for both the cytoplasmic domain (SctD^C^) and the periplasmic domain (SctD^P^), there are no high resolution structures available for any members of the SctK family [5]. SctK proteins from well-studied T3SS, such as that from *Shigella*, have proven difficult to purify for subsequent structural characterization, however, we have been able to purify sufficient quantities of the SctK protein from *P. aeruginosa*, PscK, for structure determination. We therefore crystalized and solved the structure of PscK by X-ray diffraction to a resolution of 2.5 Å (**Table 2**).

**Table 2.**
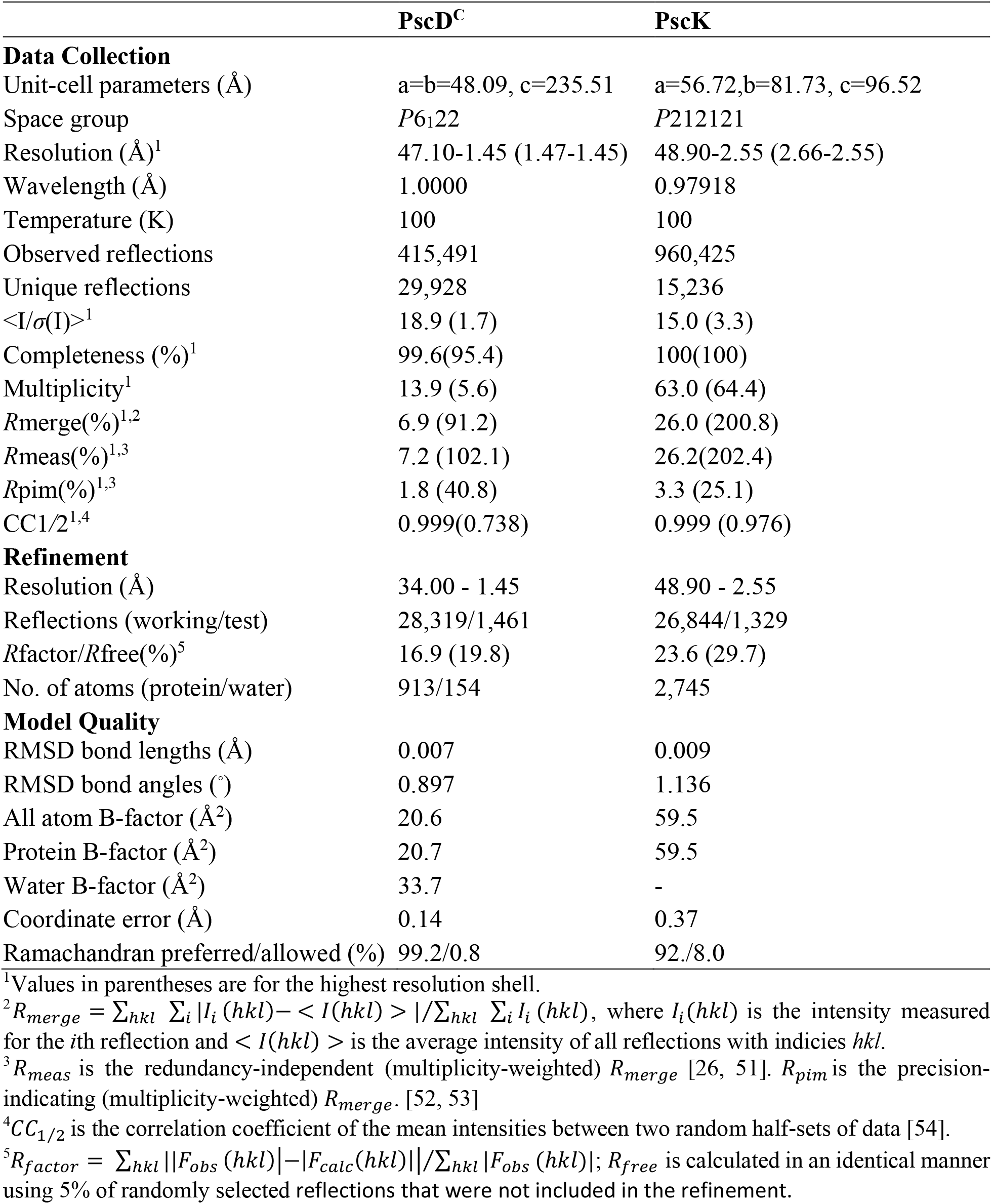
Data collection of refinement statistics.

|  | **PscD<sup>C</sup>** | **PscK** |
| --- | --- | --- |
| **Data Collection** |  |  |
| Unit-cell parameters (Å) | a=b=48.09, c=235.51 | a=56.72,b=81.73, c=96.52 |
| Space group | *P*6<sub>1</sub>22 | *P*212121 |
| Resolution (Å)<sup>1</sup> | 47.10-1.45 (1.47-1.45) | 48.90-2.55 (2.66-2.55) |
| Wavelength (Å) | 1.0000 | 0.97918 |
| Temperature (K) | 100 | 100 |
| Observed reflections | 415,491 | 960,425 |
| Unique reflections | 29,928 | 15,236 |
| $\langle I/\sigma(I) \rangle$<sup>1</sup> | 18.9 (1.7) | 15.0 (3.3) |
| Completeness (%)<sup>1</sup> | 99.6(95.4) | 100(100) |
| Multiplicity<sup>1</sup> | 13.9 (5.6) | 63.0 (64.4) |
| $R_{merge}(\%)$<sup>1,2</sup> | 6.9 (91.2) | 26.0 (200.8) |
| $R_{meas}(\%)$<sup>1,3</sup> | 7.2 (102.1) | 26.2(202.4) |
| $R_{pim}(\%)$<sup>1,3</sup> | 1.8 (40.8) | 3.3 (25.1) |
| $CC1/2$<sup>1,4</sup> | 0.999(0.738) | 0.999 (0.976) |
| **Refinement** |  |  |
| Resolution (Å) | 34.00 - 1.45 | 48.90 - 2.55 |
| Reflections (working/test) | 28,319/1,461 | 26,844/1,329 |
| $R_{factor}/R_{free}(\%)$<sup>5</sup> | 16.9 (19.8) | 23.6 (29.7) |
| No. of atoms (protein/water) | 913/154 | 2,745 |
| **Model Quality** |  |  |
| RMSD bond lengths (Å) | 0.007 | 0.009 |
| RMSD bond angles (°) | 0.897 | 1.136 |
| All atom B-factor (Å<sup>2</sup>) | 20.6 | 59.5 |
| Protein B-factor (Å<sup>2</sup>) | 20.7 | 59.5 |
| Water B-factor (Å<sup>2</sup>) | 33.7 | - |
| Coordinate error (Å) | 0.14 | 0.37 |
| Ramachandran preferred/allowed (%) | 99.2/0.8 | 92./8.0 |

<sup>1</sup>Values in parentheses are for the highest resolution shell.
<sup>2</sup> $R_{merge} = \sum_{hkl} \sum_i |I_i(hkl) - \langle I(hkl) \rangle| / \sum_{hkl} \sum_i I_i(hkl)$ , where $I_i(hkl)$ is the intensity measured for the $i$ th reflection and $\langle I(hkl) \rangle$ is the average intensity of all reflections with indices $hkl$ .
<sup>3</sup> $R_{meas}$ is the redundancy-independent (multiplicity-weighted) $R_{merge}$ [26, 51]. $R_{pim}$ is the precision-indicating (multiplicity-weighted) $R_{merge}$ . [52, 53]
<sup>4</sup> $CC_{1/2}$ is the correlation coefficient of the mean intensities between two random half-sets of data [54].
<sup>5</sup> $R_{factor} = \sum_{hkl} ||F_{obs}(hkl)| - |F_{calc}(hkl)|| / \sum_{hkl} |F_{obs}(hkl)|$ ; $R_{free}$ is calculated in an identical manner using 5% of randomly selected reflections that were not included in the refinement.

The PscK unit cell contains two molecules in the asymmetric unit, related by pseudo-translational symmetry (**Suppl. Fig. S2A**). Although PscK contains three methionine residues, only two (M1 and M57) from each subunit contributed to the anomalous signal used for structure solution. Residue M172 was in a disordered region of the polypeptide. Overall, the structures of each individual subunit are similar with an RMSD deviation between C_*α*_ atoms of 0.87 Å (175 residues) (**Suppl. Fig. S2B**). PscK adopts a fold that is composed of nine *α*-helices and three three-residue 3_10_ helices (**Fig. 1A**). The *α*-helices span the following residue ranges - *α*1: A5-F10, *α*2: A23-L28, *α*3: S39-Q48, *α*4: Q69-L82, *α*5: H84-R89, *α*6: E106-L113, *α*7: Y139-C152, *α*8: C157-R164, *α*9: Q183-A199. Portions of the loops between *α*2-*α*3, *α*5-*α*6, *α*6-*α*7, and *α*8-*α*9 are not present in the density map, possibly due to high flexibility. The largest region of missing density is 11 consecutive residues in the *α*5-*α*6 loop.

**Figure 1.**
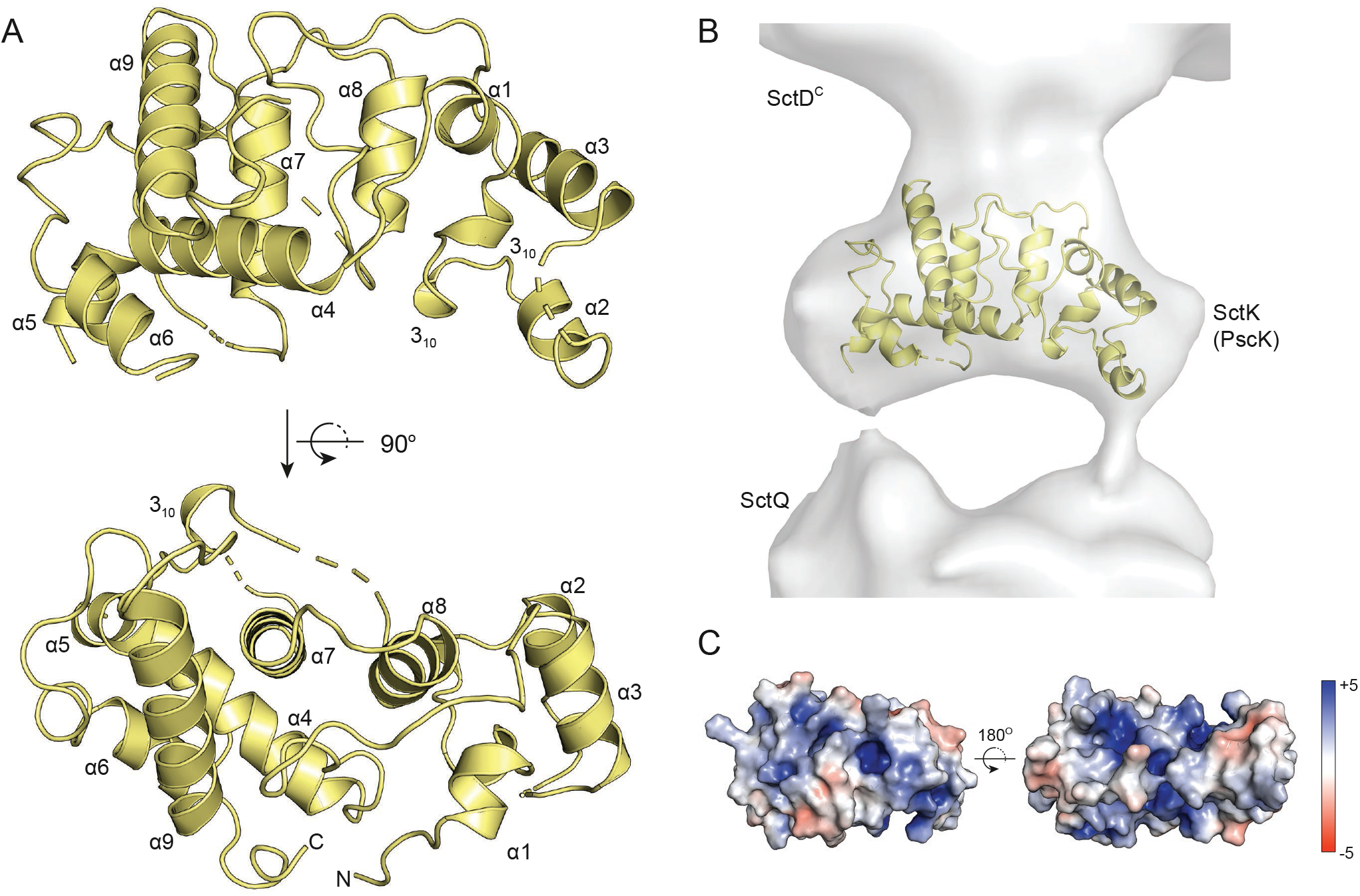
The crystal structure of PscK. **Panel A** shows a cartoon representation of the structure of PscK, with the lower depiction being rotated 90° counter-clockwise about the horizontal axis. Short regions of missing density are indicated by dashed lines. The missing segment between *α*5 and *α*6 (residues 92-102) is rendered as a gap. **Panel B** shows the PscK structure manually positioned inside the SctK EM density of the *Salmonella* injectisome. It is roughly in the same orientation as shown in the top portion of **Panel A**. The structure fits equally well when rotated 180° about the vertical axis. The solvent-excluded electrostatic surface of PscK is shown in **Panel C**, with the color gradient scaled in units of *k*_*β*_*T/eV*. With PscK positioned in the injectisome as shown in **Panel B**, the surface on the left faces SctD^C^, while the surface on the right faces SctQ.

The *α*-helical model of PscK determined from X-ray diffraction is consistent with the results of far-UV CD spectroscopy of the protein in solution. The CD spectrum of PscK shows the double minima at 222 nm and 208 nm that are characteristic of *α*-helical proteins (**Suppl. Fig. S3A**). The helicity of PscK computed from the CD spectrum is 34 %, which is close to the 42 % of residues in the X-ray structure assigned to helices. A CD thermal melt of PscK shows an unfolding transition at approximately 40 °C (**Suppl. Fig. S3B**), suggesting modest thermal stability unlike the *β*-strand rich cytoplasmic domain of proteins from the MxiG/PrgH family that melt at relatively higher temperatures [10]. This lability may explain the difficulty of purifying other members of the SctK family for structural analysis.

The kidney-shaped PscK molecule is also comparable in size and shape to the electron density of SctK molecules seen in EM studies of the T3SS [6, 9]. Manually fitting the PscK structure into the SctK density from the *Salmonella* injectisome (**Fig. 1B**) shows that PscK likely adopts an orientation in the sorting platform similar to that shown in the top panel of **Figure 1A**, with helices *α*7 and *α*8 roughly parallel to the long axis of the injectisome and with loops *α*3-*α*4 and *α*7-*α*8 facing the SctD^C^protein. PscK fits the SctK density in this orientation with the N-and C-termini pointed either outside away from the sorting platform core or rotated 180^*°*^ so that the termini face in toward the ATPase. Cryo-electron tomographic imaging of the *in situ Shigella* T3SS with bacteriophage T4 lysozyme fused at the MxiK N-terminus suggests that the termini actually face inward toward the ATPase [9].

The structure of PscK does not resemble any other known protein fold. In an effort to identify structural homologs of PscK, structure similarity searches of the PDB database were conducted using the DALI [11], PDBeFold [12], and VAST [13] servers. None of these searches found similar structures. The best match found by the DALI server was the chaperone protein DnaK from *E. coli* (PDB 5NRO) with a Z-score of 4.3 and RMSD of 3.7 Å for only 83 aligned residues. The PDBeFold server produced a single match with a very low quality score and a p-value of 1. The best VAST server result aligned only 26 residues. These results suggest that PscK represents a novel fold.

### Structure of the PscD cytoplasmic domain

It has been shown in EM structures of the *Shigella* and *Salmonella* injectisome that the cytoplasmic domain of the SctD protein forms a critical interface between the basal body IR and the sorting platform component SctK [6, 7, 9]. The structures of SctD^C^ domains from *Shigella, Yersinia*, and *Salmonella* have been solved and found to have very similar forkhead-associated (FHA) domain folds [14]. Unlike canonical FHA domains, however, the residues involved in phosphothreonine binding are either not present or at incorrect positions in these SctD^C^ FHA domains. As with SctK, there is not a great deal of sequence similarity between the SctD^C^ domains from different organisms, however, it is likely that they all adopt a similar FHA fold [10, 14, 15]. To confirm the FHA-domain structure of PscD^C^, we solved its crystal structure to a resolution of 1.45 Å (**Table 2**).

As expected, PscD^C^ forms an FHA-like domain (**Fig. 2**) that is similar to that reported for other SctD^C^ proteins (**Supp. Fig. S4**). The FHA domain signature is typically 80-100 amino acids that form an 11-strand *β*-sandwich. The PscD^C^ *β*-sandwich is formed by 11 *β*-strands that span the following residue ranges; *β*1: A2-F7, *β*2: A15-L19, *β*3: G22-G27, *β*4: D34-V36, *β*5: V45-D52, *β*6: G55-W61, *β*7: R68-Q69, *β*8: Q72-A73, *β*9: A78-I79, *β*10: Q86-C88, *β*11: L91-C96. The PscD^C^ structure is similar to high resolution structures of MxiG^C^ from *Shigella* (PDB ID: 2XXS and 4A4Y), PrgH^C^ from *Salmonella* (4G2S), and YscD^C^ from *Yersinia* (4A0E). It aligns best (RMSD = 0.56 Å) with YscD^C^ with which it also shares the highest sequence homology (**Suppl. Fig. S4**).

**Figure 2.**
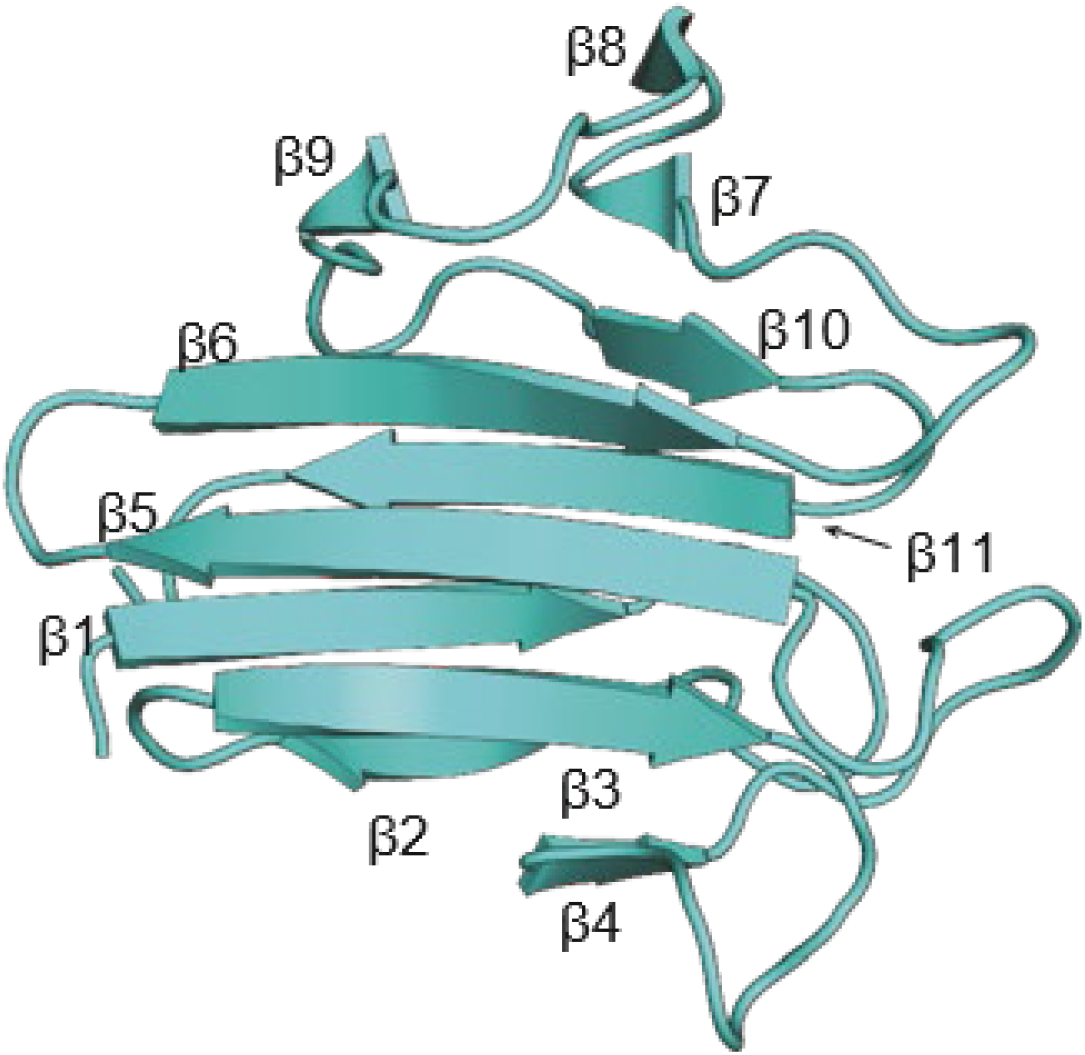
The crystal structure of PscD^C^. A cartoon representation of the PscD^C^ backbone is shown here. The *β*-sheets are labeled as described in the text. The apical face typically involved in FHA phosphopeptide binding (and phosphate independent protein-protein interactions) is formed by the loops on the right hand side of this structure.

Like all other SctD^C^ domains with known structures, PscD^C^ does not possess the conserved phosphothreonine binding motif found in canonical FHA domains [14-16]. Phosphothreonine binding in FHA domains involves two conserved residues, an arginine in the *β*3-*β*4 loop and a serine in the loop between *β*4-*β*5 [17]. PscD^C^ does not have an arginine in either of these loops and has no serine in loop *β*4-*β*5. It therefore seems likely that PscD^C^ and all other SctD^C^ domains interact with SctK proteins in a phosphothreonine-independent manner.

### Model of the PscK/PscD^C^ complex

As SctK and SctD^C^ proteins, PscK and PscD^C^ are thought to interact to connect the injectisome sorting platform to the basal body via the IR. Unfortunately, the PscK/PscD^C^ complex was not successfully crystallized, so the precise structure of the PscK/PscD^C^ complex has not yet been determined. From the structures of the individual proteins we can, however, make reasonable assumptions about the identity of the interaction surfaces of each protein and then model the PscK/PscD^C^ complex by computationally docking the proteins at those surfaces. As discussed above, the PscK molecule fits the EM density in an orientation that places the surface defined by the *α*3-*α*4 loop, the *α*7-*α*8 loop, and the loop between the third 3_10_ helix and *α*9 near the SctD^C^ density. As shown in **Figure 1C**, this surface has regions of highly positive electrostatic potential. The EM density for SctD^C^ proteins is not defined enough to definitively orient PscD^C^ within the injectisome, however, FHA domains typically interact with their ligands via the apical surface defined by loops *β*1-*β*2, *β*4-*β*5, *β*6-*β*7, and *β*10-*β*11 (**Fig. 2**). In contrast to PscK, this surface of PscD^C^ has a negative electrostatic potential and several solvent-accessible, negatively-charged residues (**Suppl. Fig. S5**).

To model the PscK-PscD^C^ interaction, the individual structures were docked together at the surfaces described above using the HADDOCK2.2 webserver [18]. HADDOCK docking converged to a single low energy docked conformation. A plot of the RMSD of the interfacial residues, using the lowest energy conformation as the reference, against the HADDOCK score for each of the 200 water-refined models generated, shows a docking funnel (**Suppl. Fig. S6**). The models with the lowest HADDOCK energies are nearly identical, with less than 0.5 Å interfacial RMSD. A representative model is shown in **Figure 3A**.

**Figure 3.**
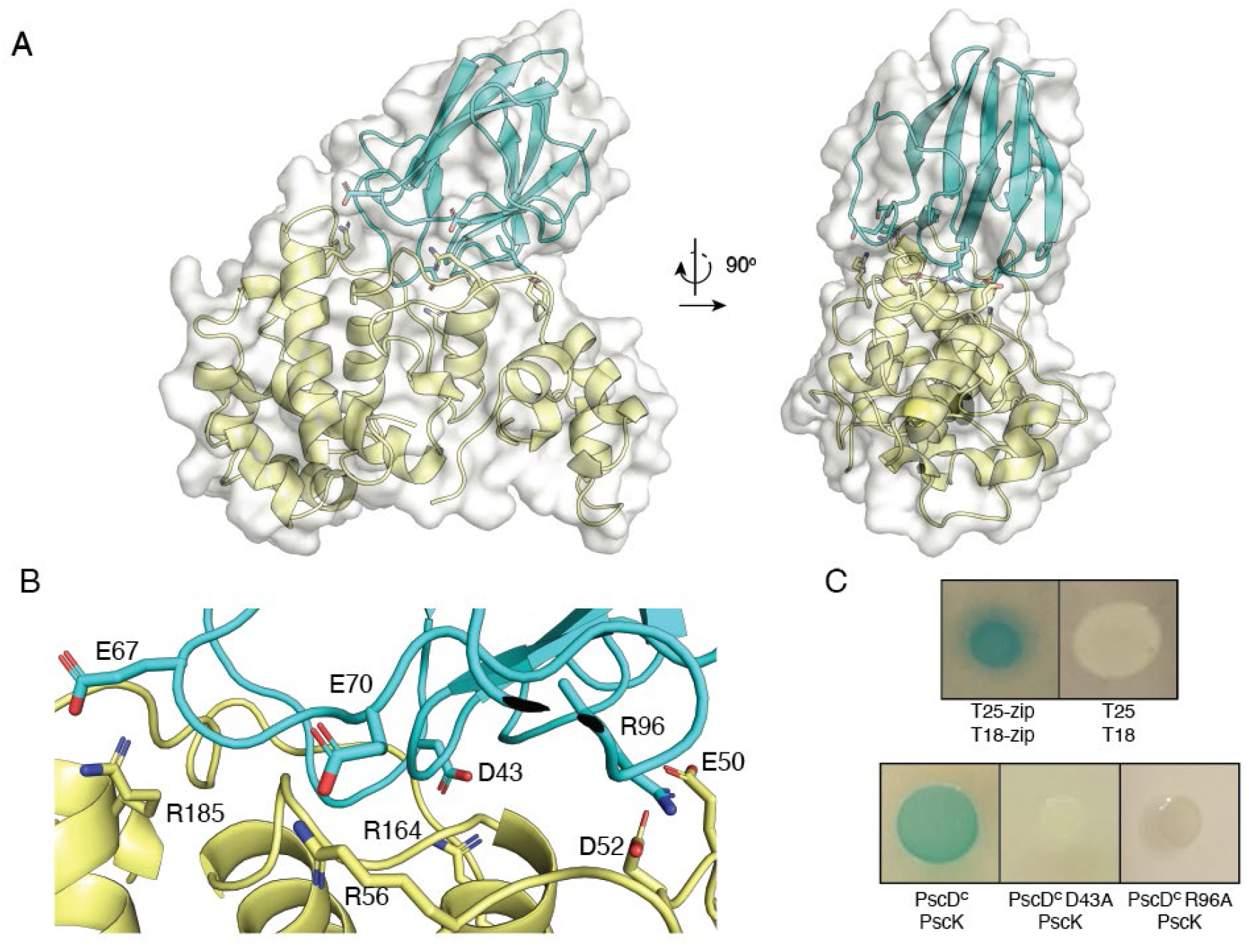
Computational docking of PscK with PscD^C^ at likely interaction surfaces converges to a single low-energy model. **Panel A** shows the entire docked complex of PscK (yellow) with PscD^C^ (cyan). A close-up of the PscK-PscD^C^ interface is shown in **Panel B**. Residues that are predicted to make important electrostatic interactions are labeled. **Panel C** shows results of bacterial two-hybrid screening for the PscK-PscD^C^ interaction. The top row shows a positive control (a leucine zipper domain fused to both the T25 and T18 AC fragments) on the left and a negative control (the T25 and T18 domains on their own) on the right. The bottom row shows results for wild-type PscK with wild-type PscD^C^ on the left, PscD^C-D43A^ in the middle, and PscD^C-R96A^ on the right. In each case, PscK was expressed with an N-terminal T18 fragment and PscD^C^ was expressed with a C-terminal T25 fragment.

Computational alanine scanning [19] of the docked PscK-PscD^C^ interface finds that the largest energetic contributions to the interaction come from charged sidechains. Interfacial residues with large ΔΔG values are D43, R96, E70, and E67 in PscD^C^ and R164, E50, R56, and R185 in PscK. As shown in **Figure 3B**, these residues appear to interact with each other across the protein-protein interface. D43, E67, and E70 of PscD^C^, respectively, form salt bridges with R164, R185, and R56 of PscK. PscD^C^ R96 is proposed to interact with two PscK residues, E50 and D52.

### In vitro testing of how mutations introduced into PscD^C^ affect the PscD^C^-PscK interaction

To test the model presented in **Figure 3**, we examined the interaction of PscK and PscD^C^ using a bacterial adenylate cylcase two-hybrid (BACTH) assay. In this experiment, each member of a protein pair is genetically fused to a fragment of the adenylate cyclase (AC) toxin from *Bordetella pertussis* (CyaA) and transformed together into an *E. coli* strain lacking a functional AC gene. Interaction between the protein pair reconstitutes the AC enzyme, which produces cAMP to induce expression of *β*-galactosidase. Successful protein-protein interactions are then detected as blue colonies on agar media plates containing X-gal.

Because the spatial configuration of the interacting proteins determines whether the T25 and T18 AC fragments will be oriented correctly to successfully reconstitute enzyme activity, we initially tested all possible combinations of wild-type PscK and PscD^C^ proteins fused with either T25 or T18 as N-or C-terminal fusions. Half of these combinations could support reconstitution of AC activity (**Fig. 3C** and **Suppl. Fig. S7**). To test the HADDOCK model of the PscK/PscD^C^ complex, we made alanine mutants of selected charged interfacial residues in the PscD^C^ domain of the PscD^C^-T25 fusion and screened for binding to T18-PscK in the BACTH system. Consistent with the model, we find that mutation of D43 in PscD^C^ appears to abolish the PscK-PscD^C^ interaction in the two-hybrid assay (**Fig. 3C**). Likewise, mutation of a prominent protruding arginine at position 96 that is modeled to interact electrostatically with two PscK residues (D52 and E50) eliminated the PscD^C^-PscK interaction in the two-hybrid assay (**Fig. 3C**), as did mutation of E70 (not shown).

Elimination of the PscD^C^ interaction with PscK could be indicative of the importance of electrostatic interactions between the proteins and the role of these residues in that process, however, it is also possible that these mutations could be compromising the folding of the PscD^C^ FHA domain. To ensure that proper folding is occurring, we used circular dichroism (CD) spectroscopy to determine whether the mutations caused any significant changes in the PscD^C^ secondary content (**Fig. 4**). The CD spectrum of PscD^C^ is characteristic for an FHA β-structure domain and this does not differ appreciably for any of the point mutants. Furthermore, when the CD signal was followed as a function of temperature, the thermal stability was not substantially different for any of these forms of PscD^C^ (**Suppl. Fig. S8**). All of these recombinant proteins displayed an onset of thermal unfolding between about 45 °C and 52 °C with the wild-type PscD^C^ having an onset of unfolding at 50 °C.

**Figure 4.**
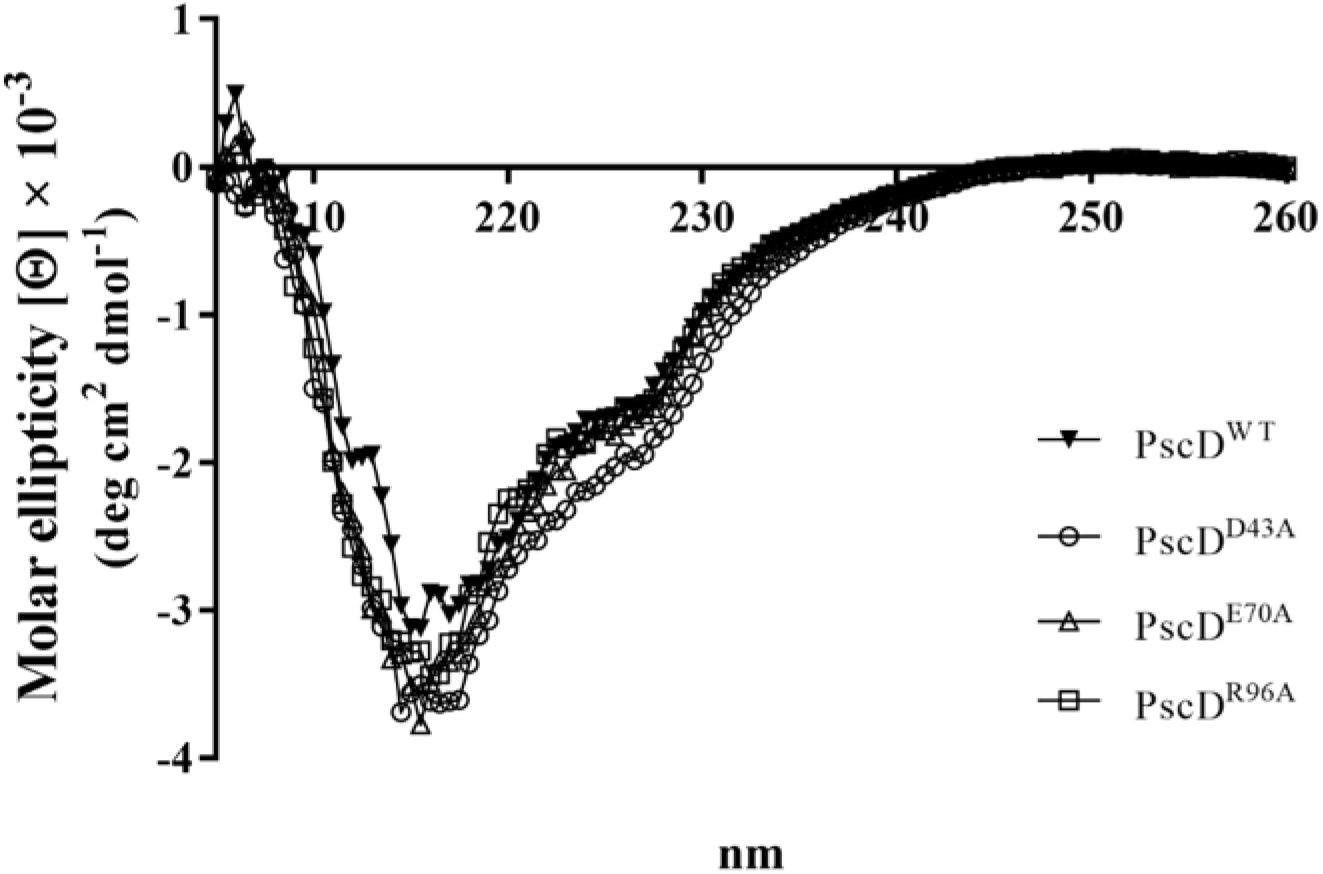
Point mutations in PscD^C^ do not alter its secondary structure profile. The wild-type PscD^C^ and each of three point mutants were purified and their CD spectra obtained. All displayed the strong β-strand component typical of FHA domains, suggesting no misfolding was occurring. Each was also found to undergo similar unfolding transitions as the temperature was increased (see **Suppl. Fig. S8**).

### Analysis of how the PscD^C^ mutations influence PscD activity in vivo

To determine whether loss of interactions in a two-hybrid analysis translate into reduced T3SS activity *in vivo*, we complemented a *pscD* null strain of *P. aeruginosa* PA14 with a chromosome integrated copy of full length wild-type *pscD* or the full length genes encoding PscD^D43A^, PscD^E70A^ and PscD^R96A^. These strains were then grown overnight at 37 °C under low-calcium conditions to induce T3SS expression and activation. The activity of the T3SS was then assessed by quantifying the phospholipase activity of the effector protein ExoU in culture supernatants [20, 21]. Because the BACTH analysis presumably measures interactions between protein pairs, it is possible that the 24-fold symmetry of the PscD ring gives rise to a high enough localized concentration of PscD^C^ to make up for the absence of a presumably pairwise PscD^C^-PscK interaction seen in the two-hybrid analysis. Likewise, it is possible that loss of a single electrostatic interaction could be offset by the existence of multiple other interactions for what is likely a transient (dynamic) association during active secretion. On the other hand, loss of T3SS activity for any of the point mutants would provide *prima facie* validation for the electrostatic-based binding model presented in **Figure 3**. As shown in **Figure 5**, while the D43A and E70A mutants appear to retain their *in vivo* functions within the context of the injectisome, the R96A mutant completely loses its ability to secrete ExoU. It thus appears that the electrostatic interactions proposed to occur between residue R96 of PscD and E50/D52 of PscK are important at the sorting platform-IR interface.

**Figure 5.**
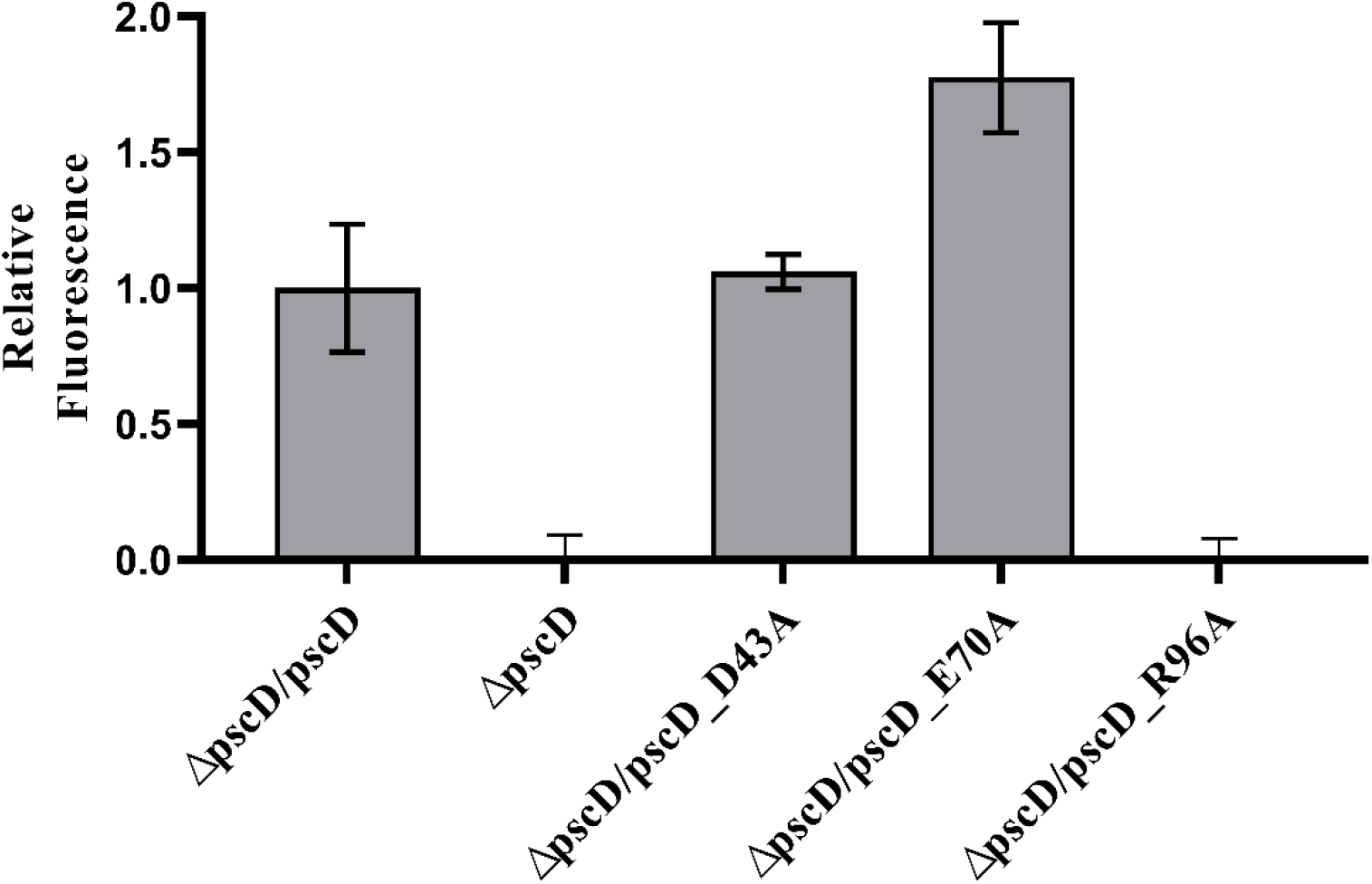
The influence of selected PscD^C^ point mutations on PscD activity *in vivo*. To determine the secretion activity of the injectisome, the relative phospholipase activity of ExoU in culture supernatants was quantified for a *P. aeruginosa* PA14 *pscD* null mutant after complementing with wild-type and mutant *pscD*. Cleavage of the PED-6 substrate resulted in an increase in fluorescence at 511 nm (excitation at 488 nm). The Δ*pscD* mutant lacking active type III secretion apparatus showed no increase in fluorescence due to the inability to secrete ExoU. Δ*pscD strain* complemented with wild-type pscD (ΔpscD/pscD) showed a much greater increase in fluorescence due to PED-6 cleavage by ExoU. The mutants *pscD*^*D43A*^ and *pscD*^*E70A*^ had activity comparable to or slightly higher than the wild-type gene, however, *pscD*^*R96A*^ had no detectable ExoU activity in the culture supernatants. Measurements represents the average of three completely independent experiments (±SE) with each performed in triplicate.

## Discussion

The overall architecture of the T3SS sorting platform appears to be generally conserved among all the pathogens possessing such systems. Nevertheless, the structures of many sorting platform components and the contributions they make to the molecular mechanisms governing type III secretion are still unknown. Recent studies have provided cryo-ET images illustrating the location of individual proteins within the sorting platform [6, 7]. In *Shigella* and *Salmonella*, the location and significance of the sorting platform SctK protein has been studied in some detail [6, 7]. SctK serves as the adaptor protein that mediates sorting platform interaction with the cytoplasmic domain of the IR protein SctD. This interaction is presumably important and the symmetry change that occurs between the basal body (SctD/MxiG/PscD with 24-fold symmetry) and the pods of the sorting platform (SctK/MxiK/PscK and SctQ/Spa33/PscQ pods with six-fold symmetry) may be indicative of dynamic interactions occurring at this interface. With regard to dynamics at this interface, it is interesting to note that SctD^C^ from *Salmonella* (PrgH^C^) clusters where it interacts with SctK (OrgA) *in situ* [6]. This is in contrast to SctD^C^ and SctK from *Shigella* (MxiG^C^ and MxiK, respectively) where the SctD^C^ domain maintains an evenly spaced 24-fold symmetry with no clustering observed [9]. This fits well with the argument that protein movements are occurring at this interface. This is also consistent with the sorting platform rearrangements in *Yersinia* observed by Diepold and colleagues [22]. Understanding this interface and the possible dynamics that could occur here will be essential to understanding the fundamental mechanisms governing type III secretion.

In the present study, we have reported the first high resolution structure for a member of the SctK family, PscK from *P. aeruginosa*, as well as the FHA domain structure of its IR interaction partner PscD. To date there are no structural or biophysical data available for any other members of the SctK protein family. Furthermore, there are no data available on any aspect of the molecular interactions that occur between the injectisome soring platform and basal body. While the *Pseudomonas* injectisome has not been structurally characterized as well as those of *Shigella* and *Salmonella*, we believe that the PscK fold is likely to be shared by the SctK proteins in those and other systems. T3SS injectisome components tend to share limited sequence conservation between species, yet they are structurally and organizationally homologous. This appears to be the case for PscK and the SctK family. Secondary structure predictions for the SctK family made with PROMALS3D [23] shows that these proteins are homologous in their distribution of secondary structure (**Suppl. Fig. S9**) despite sequence similarities ranging between 8-40% (**Suppl. Table S1**). Furthermore, the PscK structure matches the shape of the EM density for the SctK proteins in both the *Salmonella* and *Shigella* injectisomes.

We have also presented a computationally derived model of the PscK/PscD^C^ complex that connects the injectisome sorting platform to the basal body via the IR. In agreement with the calculated electrostatic surface potential of each protein, we find that the docked interface of the two proteins is energetically dependent on interactions between charged residues. In support of this model we show that for three selected positions in PscD^C^, mutation of even a single charged residue to alanine eliminates complex formation in a bacterial two-hybrid system. Whether this binding model is a general feature of SctK/SctD^C^ proteins remains to be seen. PscK is enriched with arginine residues (12.5% vs 6% natural frequency), which are often found at protein-protein interaction ‘hot spots’, and this enrichment may explain PscK’s highly positive electrostatic surface. With the exception of OrgA from *Burkholderia thailandensis*, other SctK proteins do not have significantly above average numbers of arginine residues. On the other hand, of the other SctD^C^ domains for which structures are known, YscD^C^ and MxiG^C^ have glutamate or aspartate residues at the same spatial positions as D43, E67, and E70 in PscD^C^, suggesting that this binding mode may be conserved.

To monitor the effect these single amino acid changes have on PscD activity within the context of the T3SS injectisome, a *pscD* null *P. aeruginosa* strain was complemented with either the wild-type gene or the gene harboring the point mutations D43A, E70A or R96A. These mutants failed to interact with PscK in two-hybrid assays, however, the context and possibly the presentation of the PscD^C^ FHA domain is somewhat different within the injectisome itself because it exists as a ring containing of 24 copies and in *Salmonella* it has been proposed that the PscK homologue OrgA may be interacting with multiple (perhaps four) copies of the PscD^C^ homologue PrgH^C^ (cytoplasmic FHA domain of PrgH) [6]. This is consistent with a dynamic sorting platform as proposed by Diepold *et al*. in which movement of different components has a role in regulating secretion [22]. In this model of regulation, the pod units of the SP (comprised mainly of SctQ) are in exchange between a population present within the injectisome pods and one that is a cytosolic pool of these proteins. This dynamic situation and the possibility that SctK interacts with multiple copies of the SctD FHA domain could mean that single point mutations might not be sufficient for eliminating injectisome activity. Nevertheless, one point mutation (R96A) eliminated secretion of the effector phospholipase ExoU. Meanwhile, the D43A mutation had no obvious effect on secretion and the E70A mutant actually secreted ExoU at slightly elevated levels. The fact that even one of these individual mutations eliminated ExoU secretion supports the possibility that electrostatic interactions have a role in formation of the sorting platform-IR interface with these electrostatic interactions having a role in regulating type III secretion. Future studies will fine tune this model and assist with dissection of the dynamic interactions that occur here.

## Materials and Methods

### Cloning and expression of recombinant proteins

The coding sequence for the full length *pscK* gene and that encoding the first 126 amino acids of the PscD cytosolic domain (PscD^C^) were amplified from the *P. aeruginosa* PAO1 genome. The genes were inserted into the pT7HMT expression vector so that the resulting recombinant proteins contained an N-terminal His tag that could be removed using the tobacco etch virus (TEV) protease [10]. After confirming the inserted sequences, the plasmid was used to transform *E. coli* Tuner (DE3) (Novagen). A single transformed colony was grown in LB medium with 50 µg/ml kanamycin to prepare permanent stocks for generating starter cultures.

For large-scale expression, a starter culture was diluted into fresh LB broth and grown at 37 °C with constant shaking (200 rpm) until the absorbance at 600 nm (A600) reached 0.8. Protein expression was induced by adding IPTG to 0.5 mM followed by overnight growth at 16 °C. The culture was harvested by centrifugation at 3500 × g for 30 min and the pellet was resuspended in affinity chromatography binding buffer (20 mM Tris, 0.5 M NaCl, 5 mM imidazole, pH 8.0). The cells were lysed by sonication and the lysate was clarified by centrifugation at 26,000 × g for 30 min. The protein was purified by immobilized metal affinity chromatography (IMAC) using Ni-NTA resin. The protein was eluted in binding buffer containing 500 mM imidazole and then dialyzed overnight with His-tagged TEV protease to remove the His-tag. A second IMAC column was used to remove the cleaved N-terminal tag and the TEV protease. PscD^C^ was further purified over a HiLoad 26/600 Superdex size-exclusion column equilibrated with 20 mM Tris (pH 8.0) and 200 mM NaCl. The same purification protocol was followed for PscK except that 5 % glycerol and 1 mM tris(2-carboxyethyl) phosphine (TCEP) was added to the IMAC buffers. Size exclusion chromatography was run in a suitable crystallization buffer (100 mM HEPES pH 7.5, 200 mM NaCl, 5 % glycerol and 1 mM TCEP).

Selenomethionine-(SeMet-) labeled PscK expression was expressed in an *E. coli* methionine auxotroph according to a protocol modified from Van Duyne *et al*. [24]. Cells were grown in LB broth to an A600 of 1.0. The cells were harvested as above and the pellet was resuspended in prewarmed minimal media supplemented with a standard amino acid mix (0.05 mg/mL each amino acid) in which L-Met is replaced with L-SeMet. The culture was grown at 37 °C for additional 90 min and protein expression and purification was performed as the described for native PscK.

### Crystallization PscD^C^ and PscK

Purified recombinant PscD^C^ (17.1 mg/ml in 10 mM Tris-HCl pH 7.5, 50 mM NaCl) was crystallized by vapor diffusion in Compact Jr. (Rigaku Reagents) sitting drop plates at 18 °C. After 3 days crystals grew in a SaltRx screen (Hampton Research) in well E9 (1.8 M sodium phosphate monobasic monohydrate, potassium phosphate dibasic, pH 6.9). A single prism was flash-cooled in a cryoprotectant solution consisting of 80 % mother liquor supplemented with 20 % (v/v) glycerol and diffracted to 1.45 Å resolution (**Table 2**).

PscK samples were concentrated to 20 mg/mL (native) and 10 mg/mL (SeMet) in 100 mM HEPES, pH 7.5, 200 mM NaCl, 5% glycerol, and 1 mM TCEP for crystallization screening experiments using a Formulatrix NT8 drop setting robot. Small crystals (<50 µm) of native PscK were obtained from Hampton Research SaltRx screens in condition F12. Crystals from drop F12 were used to generate a seed stock. New crystallization screens were set up with the NT8 robot using 100 nL of protein and 100 nL of crystallant and every drop was spiked with 20 nL of the seed stock. This technique produced crystals in a variety of conditions. Crystals of SeMet PscK that displayed a prismatic morphology were obtained from the Proplex HT screen (Molecular Dimensions) condition A7 (20 % (w/v) PEG 1500, 100 mM Tris, pH 7.5, 100 mM ammonium sulfate) after 1-2 days. The crystals were transferred to a fresh drop containing 80 % crystallant and 20 % (v/v) PEG before storing in liquid nitrogen. Both native and SeMet crystals grew within a week and the final structure was determined to 2.5 Å resolution (**Table 2**).

### Structure Determination, Refinement and Analysis

X-ray diffraction data for PscD^C^ (PscD residues 1-126) and PscK were collected using a Dectris Pilatus 6M pixel array detector at IMCA-CAT beamline 17ID at the APS (**Table 2**). Data were processed with XDS [25] and scaled with Aimless [26]. The PscD^C^ structure was solved by molecular replacement using MORDA, which utilized a homologous domain from the *Yersinia* protein YscD as the search model (PDB entry 4A0E). The PscK diffraction data contained a pseudotranslational symmetry peak (at 0.5, 0, 0.29) that was approximately 23 % of the origin. Attempts to solve the structure using the SAD phasing method using the SeMet data failed to yield a clear solution and produced uninterpretable electron density maps even though an appreciable anomalous signal was present as indicated from the Aimless output (DelAnom CC ∼0.54 overall). Therefore, data sets from five crystals were processed with Xia [27] using the DIALS [28] pipeline and the integrated data were scaled using Aimless, which produced a unique data set with 63-fold multiplicity (DelAnom CC ∼0.43). Following data scaling, the pseudo-translational symmetry peak had reduced to approximately 12 % of the origin. SAD phasing with Crank2 [29] using the Shelx [30], Refmac [31], Solomon [32], Parrot [33]and Buccaneer [34] pipeline via CCP4 [35] interface yielded a mean figure of merit of 0.48 following phasing/density modification. Subsequent automated model building/refinement within the Crank2 pipeline produced a model that contained 401 out of 420 possible residues and refined to R/Rfree = 30.6 %/36.8 % (space group P212121). All refinement and manual model building were conducted with PHENIX [36] and Coot [37], respectively. Disordered side chains were truncated to the point for which electron density could be observed. Structure validation was conducted with Molprobity [38] and figures were prepared using the CCP4GM package [39]. Structure superposition was carried out using GESAMT [40]. TLS refinement [41] was incorporated in the final stages to model anisotropic atomic displacement parameters for PscD^C^.

### Molecular docking of PscK and PscD^C^

The crystal structure of PscK has four loops with missing density. Those loops were modeled with RosettaCM [42], using Chain A from the crystal structure as the template. The model with the best Rosetta score was selected for docking studies. Linker residues of PscD^C^ following P111 were truncated. Restrained docking was subsequently performed using the HADDOCK2.2 webserver [18], which performs docking using ambiguous distance restraints between all residues in the protein-protein interface. Active residues (those deemed more important for the interaction) of PscK were defined as 52-62, 153-159, and 173-185. Active residues of PscD^C^ were defined as 13-18, 43-49, 67-78, and 93-96. Passive residues (those neighboring the active residues) were defined automatically by the HADDOCK server. Computational alanine scanning of the docked PscK-PscD^C^ interface was performed using Rosetta [19].

### Bacterial two-hybrid assay

The PscK-PscD^C^ interaction was tested using the bacterial adenylate-cyclase two-hybrid system (BACTH kit, Euromedex, France) [43, 44]. The coding sequences of PscK and PscD^C^ were cloned into the vectors pKT25, pKNT25, pUT18 and pUT18C as fusions with the T25 and T18 fragments using BamHI and KpnI restriction sites. The cloned constructs were verified by sequencing using T25 5’ primer (GCGCAGTTCGGTGACCAGCG) and T18 (CGGATAACAATTTCACACAG). Compatible hybrid plasmids covering all the combinations of partners for potential PscK-PscD^C^ interactions were co-transformed into *E. coli* BTH101 (lacking any adenylate cyclase). All growth media were supplemented with ampicillin and kanamycin to select for the BACTH plasmids. For X-gal detection, colonies expressing the fusion plasmid pairs were induced in LB broth with 0.5 mM IPTG overnight. Pre-induced cells then were spotted onto LB-agar indicator plates that included 40 µg /ml X-Gal with 0.5 mM IPTG. Plates were incubated at 30 °C overnight at which point positive interactions were observed as blue colonies.

### Circular dichroism (CD) spectroscopy

CD spectra were collected with 0.5 mg/ml purified protein in sodium citrate buffer (pH 7.0). Spectra were obtained in the far-UV region (190-260 nm) using a Jasco J-1500 spectropolarimeter with 0.1 cm path-length quartz cuvette at 10 °C with a spectral resolution of 0.2 nm, a scanning rate of 50 nm/min, and a 2 s integration time. Baseline corrected data was converted into molar ellipticity [*θ*] using parameters including the protein concentration, cuvette path-length (cm), molecular weight and number of amino acids. Percent helicity of PscK was calculated using the BeStSel method [45]. Thermal unfolding of the protein was determined as 222 nm over a temperature range from 10 °C to 90 °C in 0.2 °C increments with data acquired every 2.5 °C at a temperature ramp of 15°C/hour. All measurements were done in triplicate and averaged.

### Complementation of a P. aeruginosa pscD null strain using a site specific mini-Tn7 based insertion system

A *P. aeruginosa* PA14 *pscD* null strain was kindly provided by D.W. Frank [46]. Single copy complementation with wild-type or mutated *pscD* was done by inserting the gene into a conserved site on the bacterial chromosome downstream of *glmS* (glutamine-fructose-6-phosphate aminotransferase) [47]. Full length *pscD* was amplified by PCR using gene specific primer (**Suppl. Table S2**). The PCR product was ligated into BamHI-and KpnI-digested pUC18-mini-Tn7T-LAC and placed under the control of an IPTG-inducible promoter. The *pscD*-containing fusion plasmid and a pTNS2 helper plasmid were co-electroporated into the *pscD* null strain (*ΔpscD*) for which electrocompetent cells were prepared as described [48]. The cells were incubated for 1 h in LB broth and then plated on LB agar with 30 µg/mL gentamycin (Gm) and incubated for 1-2 days. To confirm the orientation of the gene insertion into a single *att* Tn7 site in the chromosome, colony PCR was performed using PglmS Up/PscD forward and pTn7R/PglmS down primer pairs and verified by sequencing. PCR-based site*-*directed mutagenesis with In-Fusion *HD* Cloning system (Takara, Clonetech) was used to generate PscD D43A, E70A and R96A mutants.

### Measurement of ExoU activity in P. aeruginosa culture supernatants

*P. aeruginosa* type III effector proteins (ExoS, ExoT, ExoY and ExoU) are actively secreted when grown in a calcium free environment. ExoU phospholipase activity in culture supernatants was quantified using the fluorogenic phospholipid analog PED-6 (*N*-((6-(2,4-dinitrophenyl) amino) hexanoyl)-2-(4,4-difluoro-5,7-dimethyl-4-bora-3a,4a-diaza-*s*-indacene-3-pentanoyl)-1-hexadecanoyl-*sn*-glycero-3-phosphoethanolamine) (Invitrogen). Cleavage of PED-6 eliminates a self-quenching effect that results in the release of a BODIPY-FL dye. This is monitored by measuring an increase in fluorescence intensity [20, 21] with the assay was performed as previously described [49, 50] in 50µl final volume containing the reaction mixture (50 mM MOPS (pH 6.3), 50 mM NaCl, K48-linked polyubiquitin (Enzo Life Sciences), 250 mM monosodium glutamate (MSG) and 30 µM PED6). Relative fluorescence intensity (RFU) was measured using a Spectramax M5 microplate reader (Molecular devices) with an excitation wavelength of 488 nm and with emission measured at 511 nm.

Briefly, *P. aeruginosa* cells complemented to express wild-type or mutant *pscD* were grown overnight at 37°C in LB broth containing 5mM EGTA and 20 mM MgCl_2_ (T3SS inducing medium). To assess the effect of the site specific mutations on ExoU secretion, *Pseudomonas* expressing wild-type *pscD* or *pscD* harboring the mutations D43A, E70A, R96A were grown in T3SS inducing medium supplemented with 30 µg/mL gentamycin and 5 mM IPTG. The *Pseudomonas* culture supernatants are then added to the reaction mixture for measuring the relative fluorescence with high levels of fluorescence indicating that ExoU has been secreted.

## Accession numbers

The coordinates and structure factors for the crystal structures generated in this study have been deposited to the Worldwide Protein Databank with the accession codes 6UID (PscD) and 6UIE (PscK).

## Acknowledgements

This research was funded by awards from the NIH (NIAID R01 AI123351 and R21 AI146517) to WDP. Use of the University of Kansas Protein Structure Laboratory was supported by a grant from the National Institutes of Health (NIGMS P30 GM110761). Use of the IMCA-CAT beamline 17-ID at the Advanced Photon Source was supported by the companies of the Industrial Macromolecular Crystallography Association (IMCA) through a contract with Hauptman-Woodward Medical Research Institute. Use of the Advanced Photon Source was supported by the U.S. Department of Energy, Office of Science, Office of Basic Energy Sciences, under Contract No. DE-AC02-06CH11357. Use of the Computational Chemical Biology Laboratory was made possible by a grant from the National Institutes of Health (NIGMS P20 GM113117). Thanks to Dr. Josephine Chandler for providing pUC18-mini-Tn7T-LAC system.

## Author contributions

W.D.P. was responsible for conceptualization, funding acquisition, project administration, writing original draft, resources, supervision; M.M. was responsible for data curation, formal analysis, investigation, methodology, reviewing and editing; S.K.W. was responsible for conceptualization, formal analysis, investigation, methodology, visualization, writing original draft; W.L.P was responsible for conceptualization and manuscript editing and writing; S.L. was responsible for data curation, formal analysis, resources, validation, visualization, review; K.P.B. was responsible for investigation, methodology, resources, software, validation; S.T. was responsible for investigation, methodology; D.K.J. was responsible for data curation, formal analysis, software, validation, visualization.

## Glossary

T3SS: type III secretion system
injectisome: type III secretion apparatus
Sct: secretion and cellular translocation
BACTH: bacterial adenylate cyclase two-hybrid
FHA domain: forkhead-associated protein domain
CD: circular dichroism

## Supplemental Materials

**Supplemental Figure S1.**
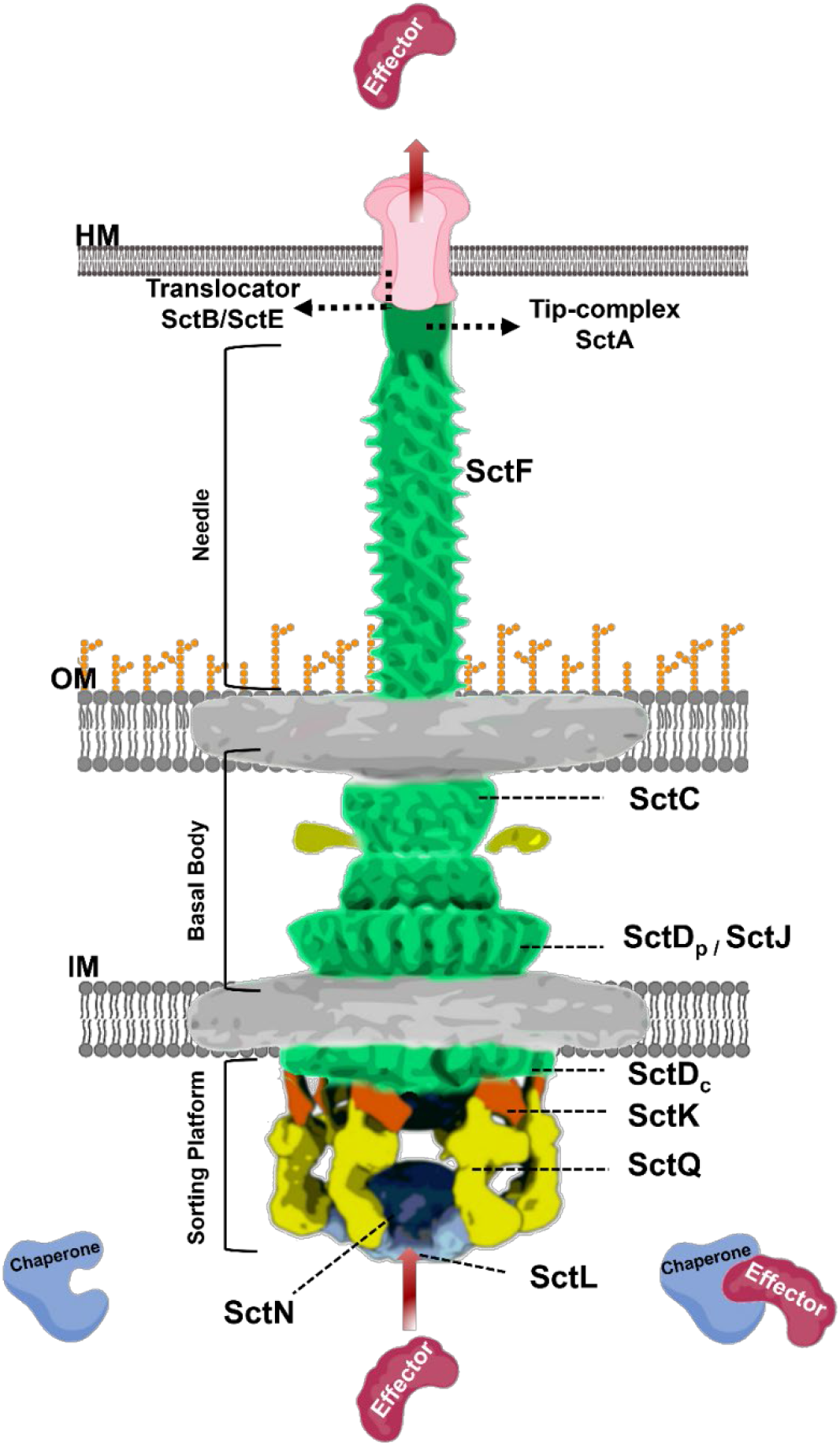
Architecture of the T3SS injectisome. The injectisome consists of an extracellular needle and tip complex that contacts the host membrane (HM), a basal body assembly that spans both the inner (IM) and outer (OM) cellular membranes, and a cytosolic sorting platform. The sorting platform SctQ proteins (yellow) interface with the basal body SctD proteins (green) via SctK adaptor proteins (orange). SctD possesses a densely packed C-terminal domain located in the bacterial periplasm (SctD_P_ or SctD^P^), a single pass transmembrane helix (not shown) and a small periplasmic domain (SctD_C_ or SctD^C^) that is connected to the transmembrane helix by an approximately 15 to 20 residue peptide linker.

**Supplemental Figure S2.**
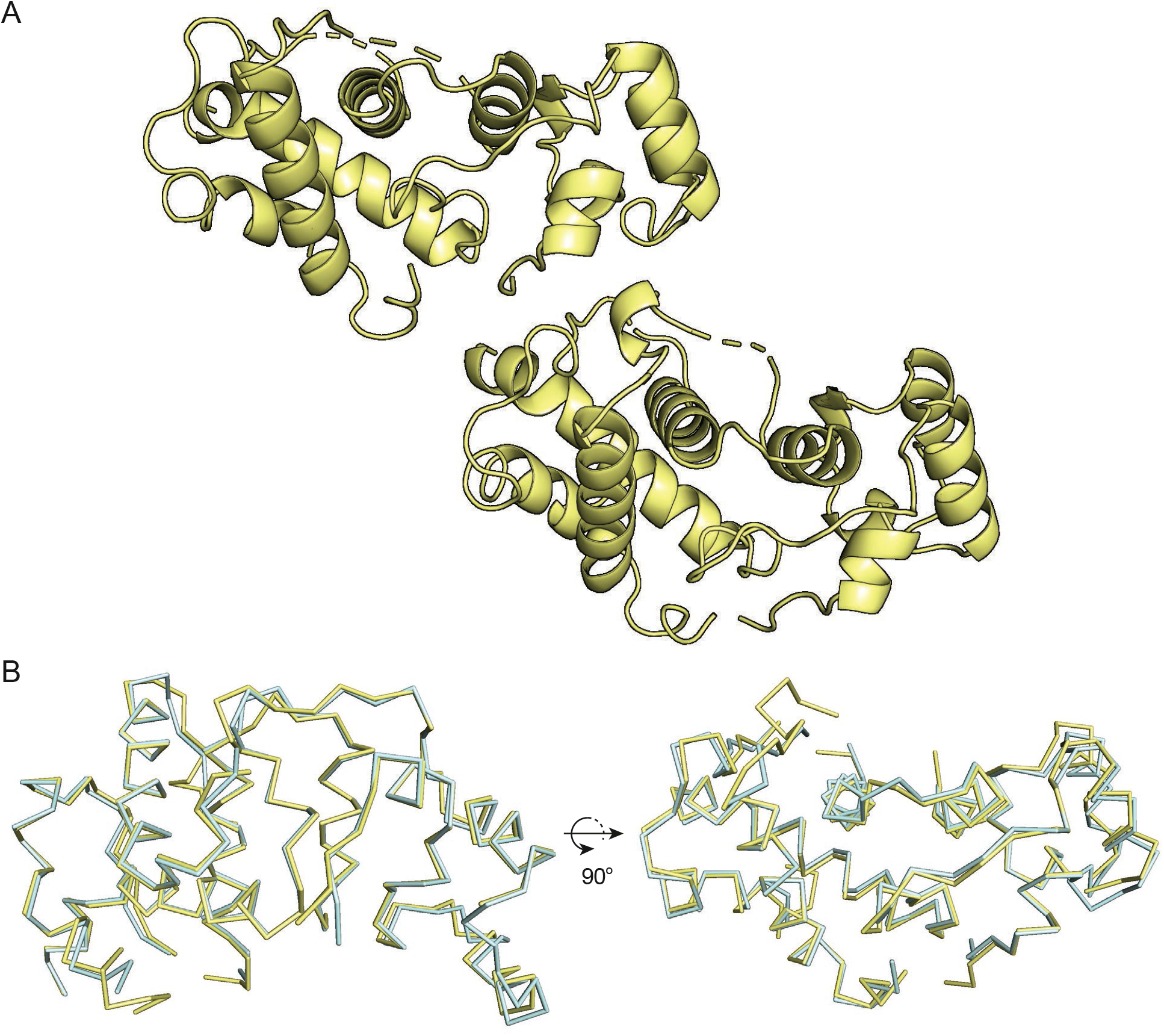
PscK molecules in the crystallographic asymmetric unit are related by a pseudo-translational symmetry as shown in **Panel A**. The models for each molecule are superimposed in **Panel B**, with one in yellow and the other in cyan.

**Supplemental Figure S3.**
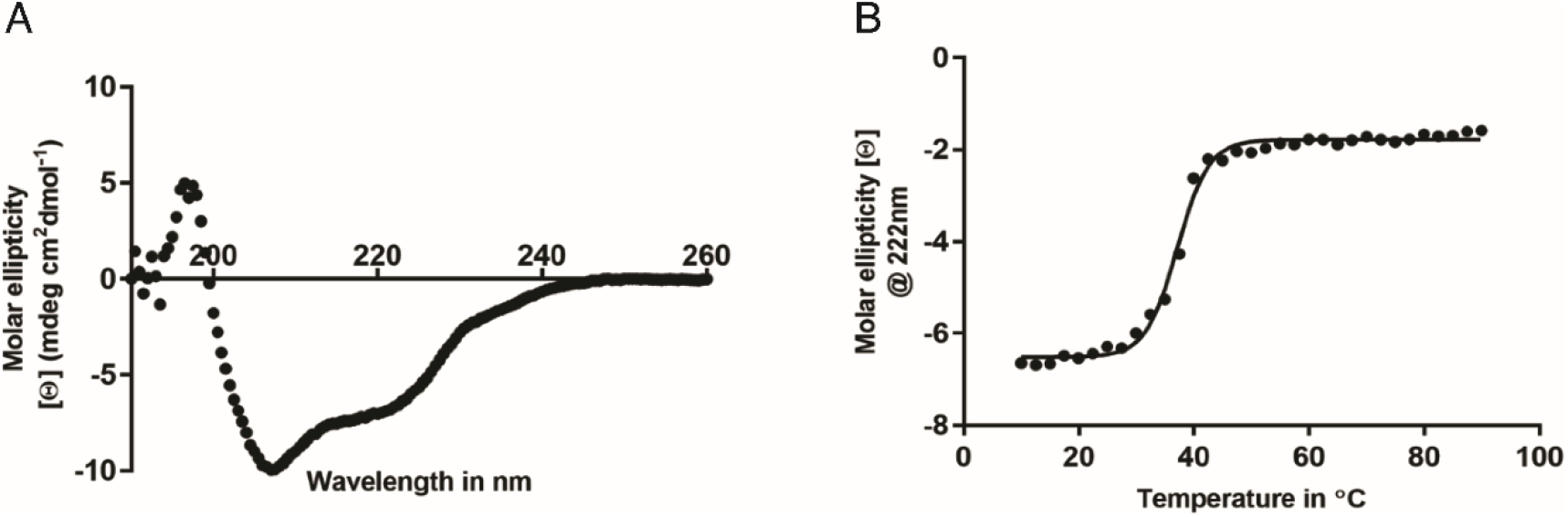
Far-UV circular dichroism (CD) spectroscopy of PscK. PscK give a CD spectrum characteristic of α-helical proteins, shown in **Panel A**, with minima at 222 nm and 208 nm. **Panel B** shows the mean residue ellipticity measured at 222 nm for PscK as the temperature is increased from 10 °C to 90 °C.

**Supplemental Figure S4.**
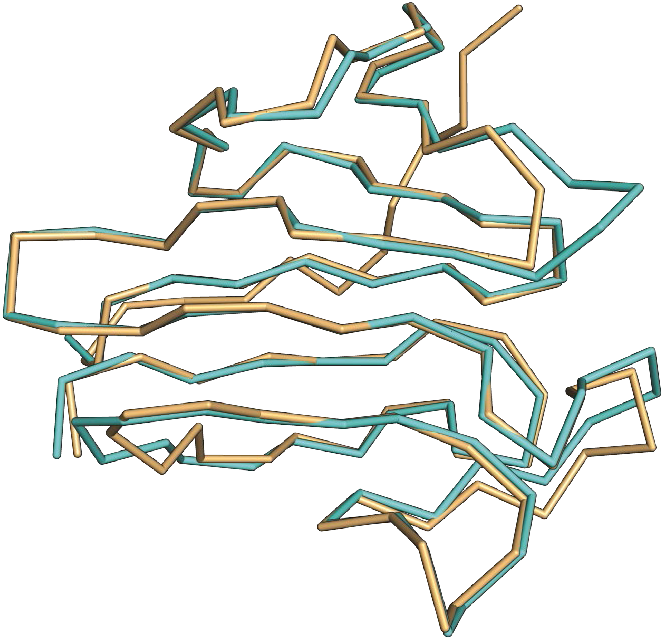
An overlay of the PscD^C^ backbone over its most closely related homologue. This is an overlay of the PscD^C^ backbone in cyan with the backbone of YscD^C^ (PDB: 4A0E) in light orange. The two structures have a backbone RMSD of 0.56 Å.

**Supplemental Figure S5.**
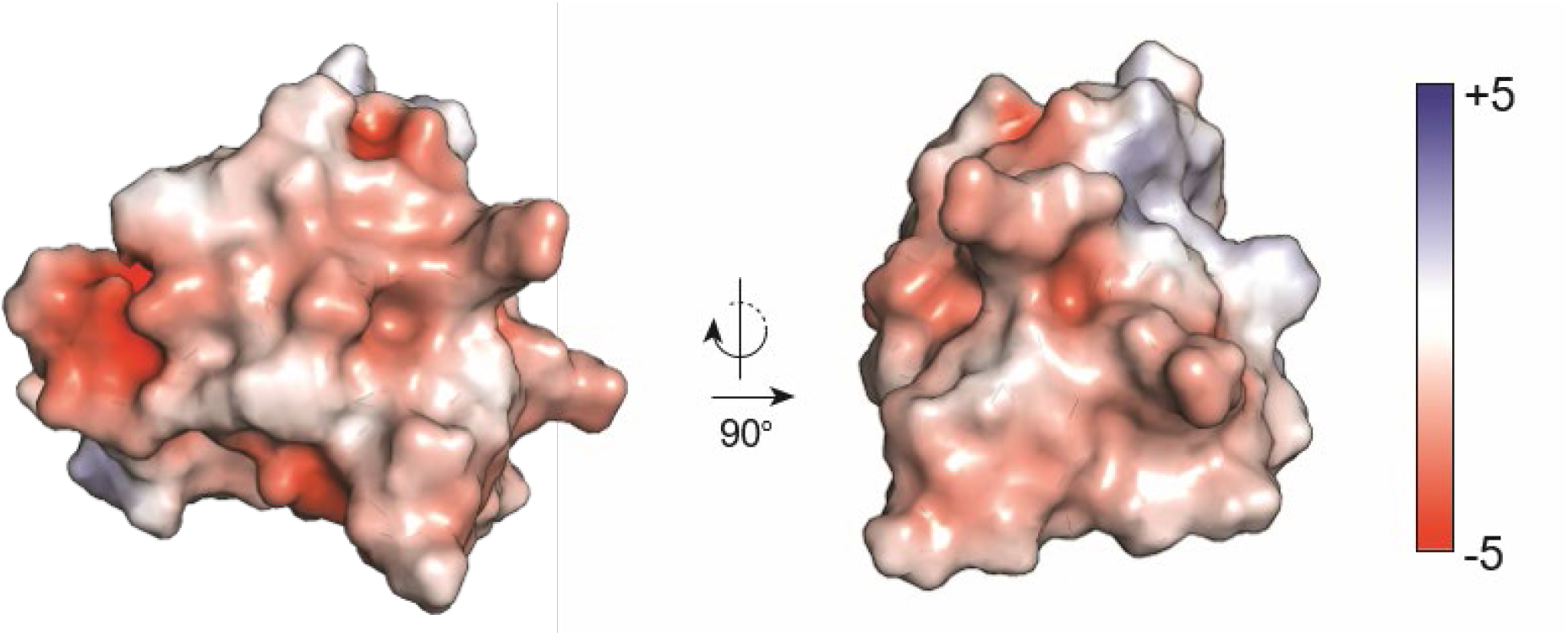
APBS electrostatic surface of PscD^C^. On the left, PscD^C^ is in the same orientation as shown in **Figure 2** of the main text. The surface of PscD^C^ that is typically involved in ligand binding in FHA domain proteins and likely faces PscK is shown on the right.

**Supplemental Figure S6.**
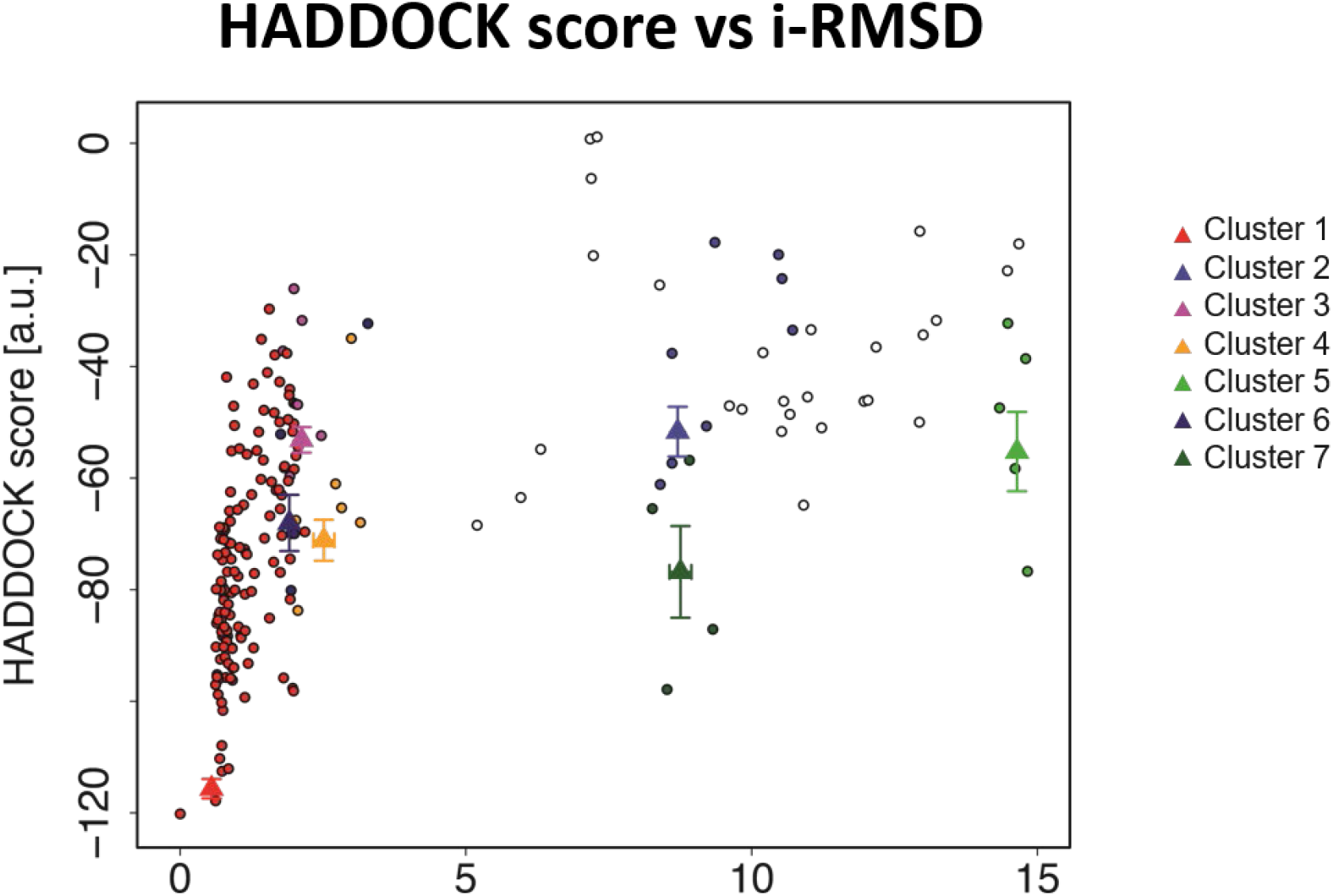
Plot of HADDOCK energy score vs. interface RMSD (i-RMSD) for all 200 water-refined models generated by the HADDOCK2.2 docking of PscK and PscD^C^. The interface i-RMSD is calculated for each model in reference to the lowest energy structure. Models fall into 7 clusters, with cluster 1 representing 69% of all models.

**Supplemental Figure S7.**
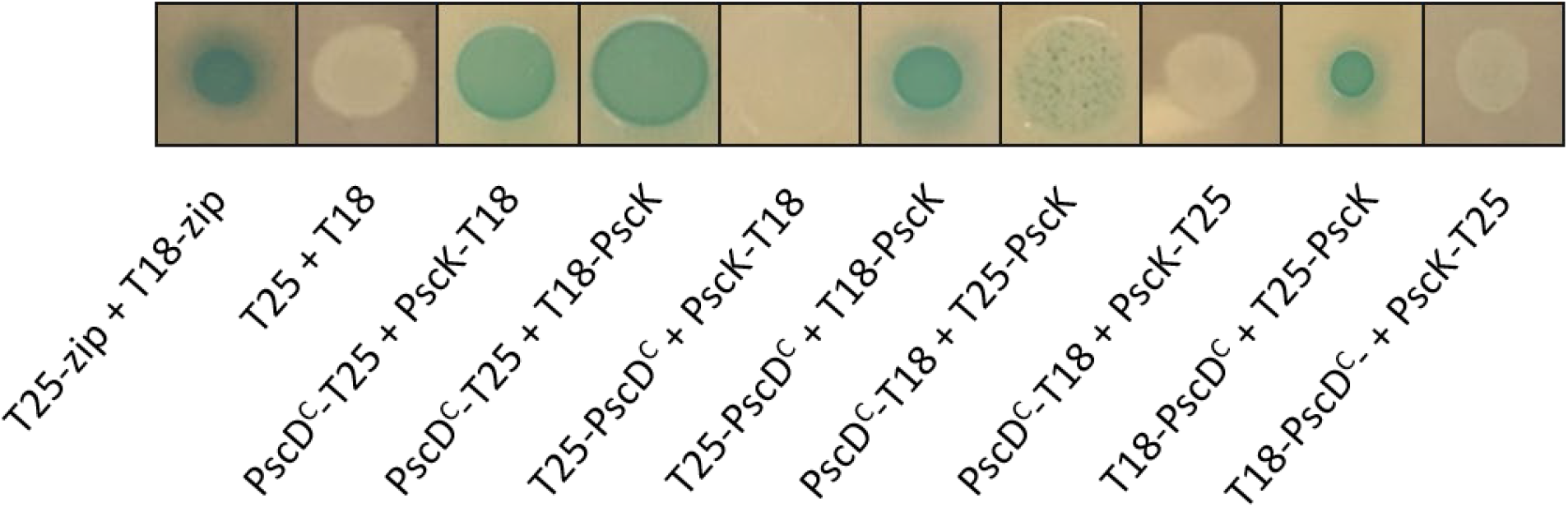
BACTH assay for PscK and PscD^C^ interaction. All possible combinations of N-and C-terminal fusions of PscK and PscD^C^ to the T25 and T18 fragments of adenylate cyclase were tested. Cells were grown in liquid media then spotted onto LB agar plates containing X-gal. A positive interaction gives blue colonies. T25-zip and T18-zip are C-terminal fusions to a leucine zipper domain and used as a positive control. The negative control is the T18 and T25 vectors lacking the bait and prey proteins.

**Supplemental Figure S8.**
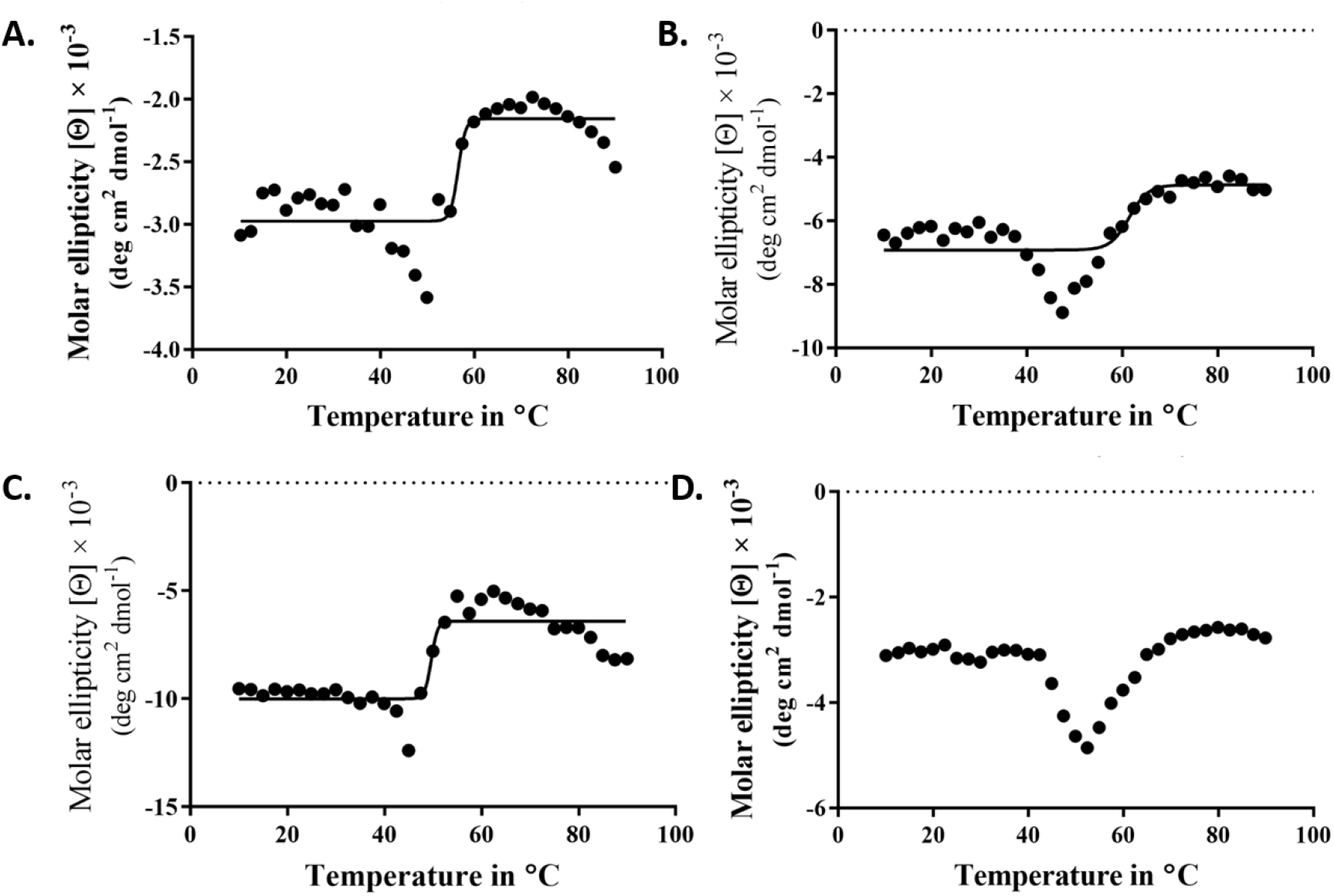
The thermal unfolding of PscD^C^ and PscD^C^ point mutants was determined by fixing the wavelength (at 218 nm) and monitoring the change in signal as a function of temperature. The onset of thermal disruption or unfolding of the secondary structure was considered the point at which the signal reached its lowest point and then started to rise. This point was approximately 50 °C for wild-type PscD^C^ (A), 48°C for PscD^C-D43A^ (B), 45 °C for PscD^C-E70A^ (C) and 52 °C for PscD^C-R96A^ (D).

**Supplemental Figure S9.**
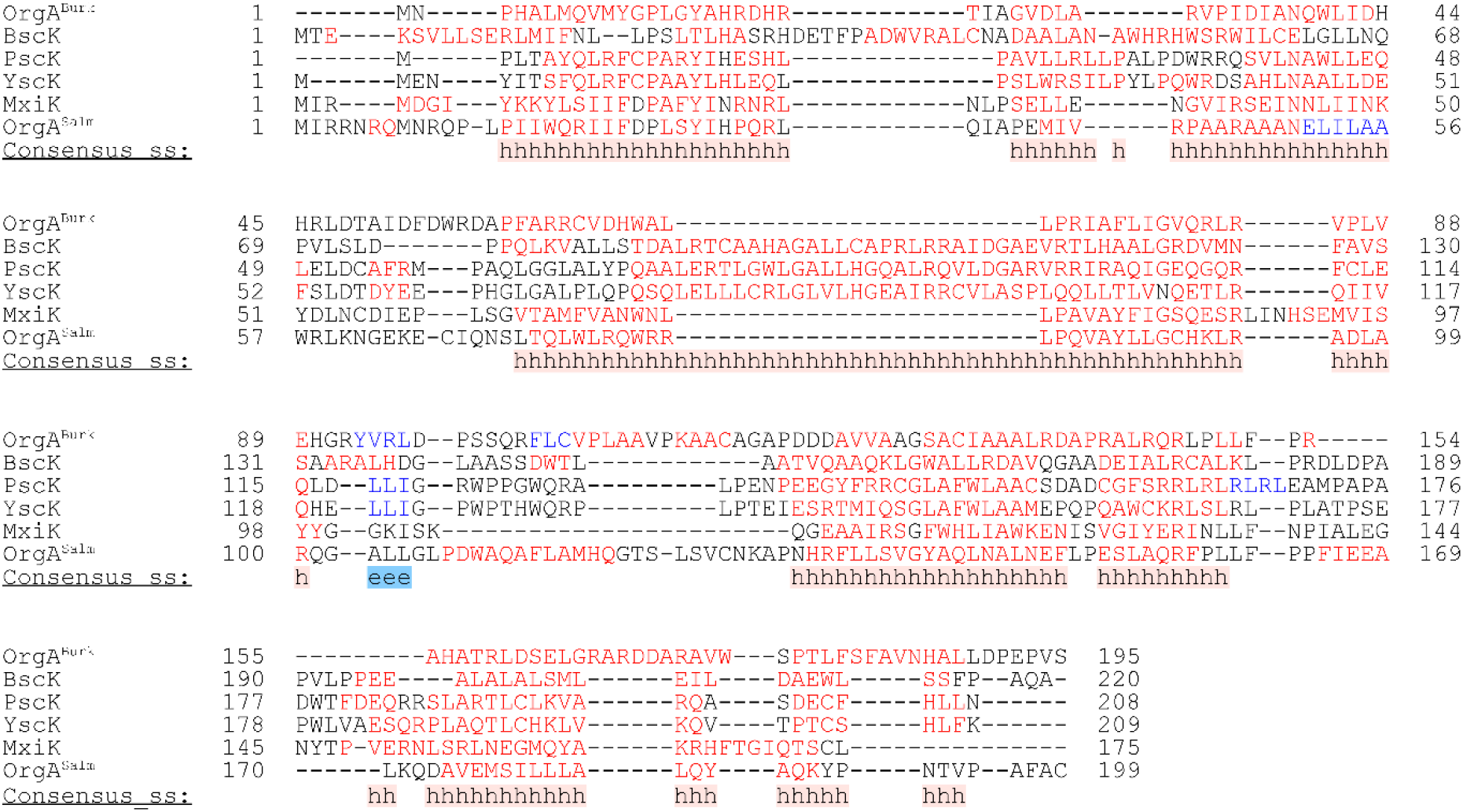
PROMALS3D multiple sequence alignment and secondary structure prediction for SctK-family proteins. Residues in helices are shown in red, those in blue are in *β*-strands. The consensus secondary structure for each position is shown at the bottom for helices (h) or sheets (e). Proteins not listed under the Sct system in Table 1 are from *Bordetella bronchiseptica* (BscK) and *Burkholderia thailandensis* T3SS-3 (OrgA^Burk^). [J. Pei, B.H. Kim, N.V. Grishin. PROMALS3D: A tool for multiple protein sequence and structure alignments. *Nucleic Acids Research* 36 (2008) 2295–2300]

**Supplemental Table S1.**
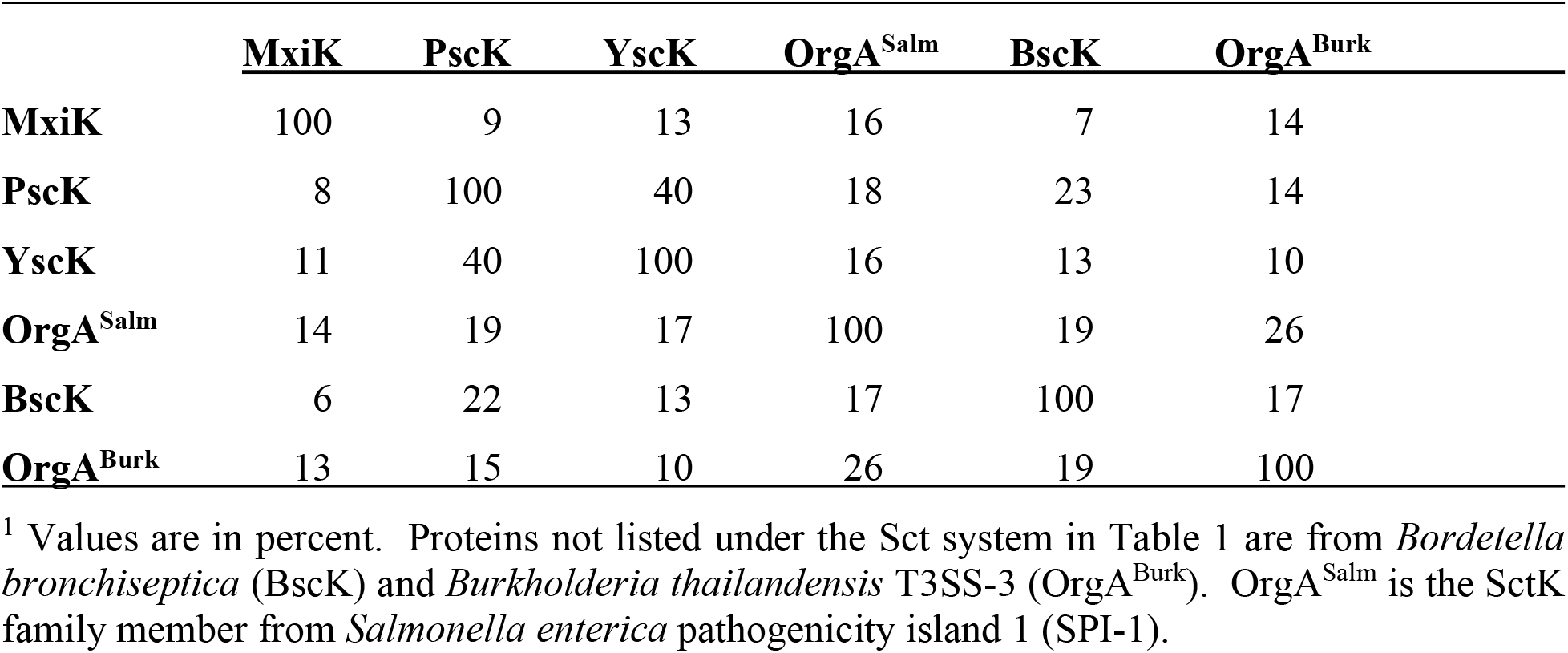
Sequence similarity matrix between SctK family proteins^1^.

**Supplemental Table S2.**
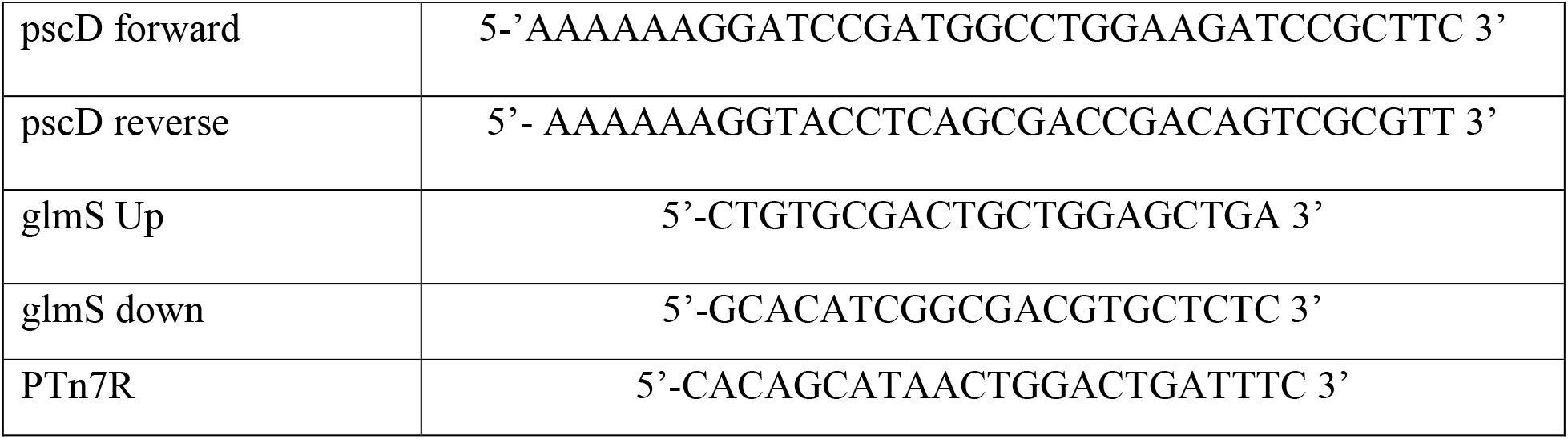
Primers used for generating constructs for complementing the *P. aeruginosa* PA14 *pscD* null strain.

